# When polymorphism and monomorphism meet: discordant genomic and phenotypic clines across a lizard contact zone

**DOI:** 10.1101/840678

**Authors:** Caroline M. Dong, Claire A. McLean, Adam Elliott, Adnan Moussalli, Devi Stuart-Fox

## Abstract

Colour polymorphism can promote rapid evolution and speciation, particularly when populations differ in the number or composition of morphs. The contact zone between *Ctenophorus modestus* (swift dragon) and *C. decresii* (tawny dragon) is a compelling study system in which to examine evolutionary processes and outcomes when polymorphic and monomorphic populations meet. *Ctenophorus modestus* is polymorphic for male throat coloration and lacks ultraviolet (UV) reflectance while *C. decresii* is monomorphic with UV-blue throats. We characterised genomic and phenotypic clines across the contact zone based on single nucleotide polymorphisms, the mitochondrial ND4 gene, and male colour traits, and concurrently assessed the phenotype of captive-bred F1 hybrids. Our results indicate that genomic introgression is asymmetric, with high frequencies of backcrossing to *C. modestus* but not *C. decresii*, accompanied by the prevalence of the *C. modestus* mtDNA haplotype in hybrids. The clines for throat phenotype are abrupt and displaced towards the range of *C. decresii*, relative to the genetic and dorsolateral phenotype clines. By contrast, both throat and dorsolateral phenotypes in captive-bred F1 hybrids are intermediate. Our results are consistent with the hypothesis that throat coloration, a polymorphic sexual signal in *C. modestus*, is the target of selection during incipient speciation and provide insight into the microevolutionary processes that may link polymorphism and speciation.

## Introduction

Colour polymorphism, the co-existence of multiple colour morphs within a single interbreeding population (Huxley, 1955), is possibly the best studied model of discrete intraspecific phenotypic variation. Recently, there has been great interest in the role of colour polymorphism in promoting rapid phenotypic evolution and speciation, particularly when populations differ in the number, type, and frequency of morphs (Corl, Davis, Kuchta, & Sinervo, 2010; Forsman, Ahnesjö, Caesar, & Karlsson, 2008; Gray & McKinnon, 2007; Hugall & Stuart-Fox, 2012; McLean & Stuart-Fox, 2014; West-Eberhard, 1986). However, we lack detailed studies on the microevolutionary processes linking colour polymorphism and speciation. In particular, the genomic and phenotypic consequences of secondary contact between polymorphic and monomorphic lineages remain unclear, especially where the monomorphic lineage represents a novel morph.

Based on the widespread importance of colour signals in reproductive isolation (Boughman, 2001; Hooper, Griffith, & Price, 2019; Jiggins, Naisbit, Coe, & Mallet, 2001; Mavárez et al., 2006; Sætre et al., 1997; Seehausen et al., 2008) and the predicted genetic architecture of polymorphism (Sinervo & Svensson, 2002), there is expected to be a high probability of incompatibility during secondary contact between populations that differ in morph number, type, or frequency. Gene flow may result in the breakdown of adaptive genetic correlations and generate genetic incompatibilities between morphs (Pryke & Griffith, 2009). More generally, incompatibilities may arise when coadapted alleles are brought together for the first time in hybrids (Dobzhansky-Muller incompatibilities; Dobzhansky, 1937; Muller, 1942) or when coadapted nuclear and organelle (e.g. mitochondria and chloroplast) genomes that evolved in allopatry are disrupted (cytonuclear incompatibilities; Burton and Barreto 2012; Hill 2015). Further, a divergence in sexual phenotypic characteristics may also be accompanied by behavioural incompatibilities in mating preferences resulting in restricted gene flow and/or selection against hybrids. These phenotypic and genetic incompatibilities together may facilitate the process of speciation between populations which differ in morph frequencies and/or composition.

Here, we investigate the outcome of contemporary contact between the colour polymorphic *Ctenophorus modestus* (swift dragon; Ahl, 1926) and monomorphic *C. decresii* (tawny dragon; Duméril & Bibron, 1837), previously regarded as divergent lineages of *C. decresii* sensu lato (Dong et al., in press). The two species differ in male dorsolateral and throat coloration (Houston 1974; McLean et al. 2014b; Figures 1 and 2a). The polymorphic *C. modestus* (previously northern lineage *C. decresii*) has four discrete male throat colour morphs that are present in all populations and reflect little to no ultraviolet (UV): orange, yellow, yellow with orange centre, and grey (Teasdale et al. 2013; Figure 1a). The black dorsolateral stripe of *C. modestus* is relatively straight-edged and continuous, sometimes with one distinct blotch near the neck, and with yellow/orange coloration terminating just behind the shoulder (Figure 2a; McLean, Moussalli, Sass, & Stuart-Fox, 2013). By contrast, monomorphic *C. decresii* (previously southern lineage) males have blue throat coloration with a strong UV reflectance peak, representing a novel morph (Figure 1b; McLean et al. 2014b). Males within an insular population (i.e. Kangaroo Island; “KI”) have yellow reticulations in addition to the UV-blue coloration (Figure 2a; McLean et al. 2014b). The black dorsolateral stripe of *C. decresii* is discontinuous with two distinct blotches near the neck and coloration terminating mid-body (Figure 2a; McLean et al., 2013).

**Figure 1.**
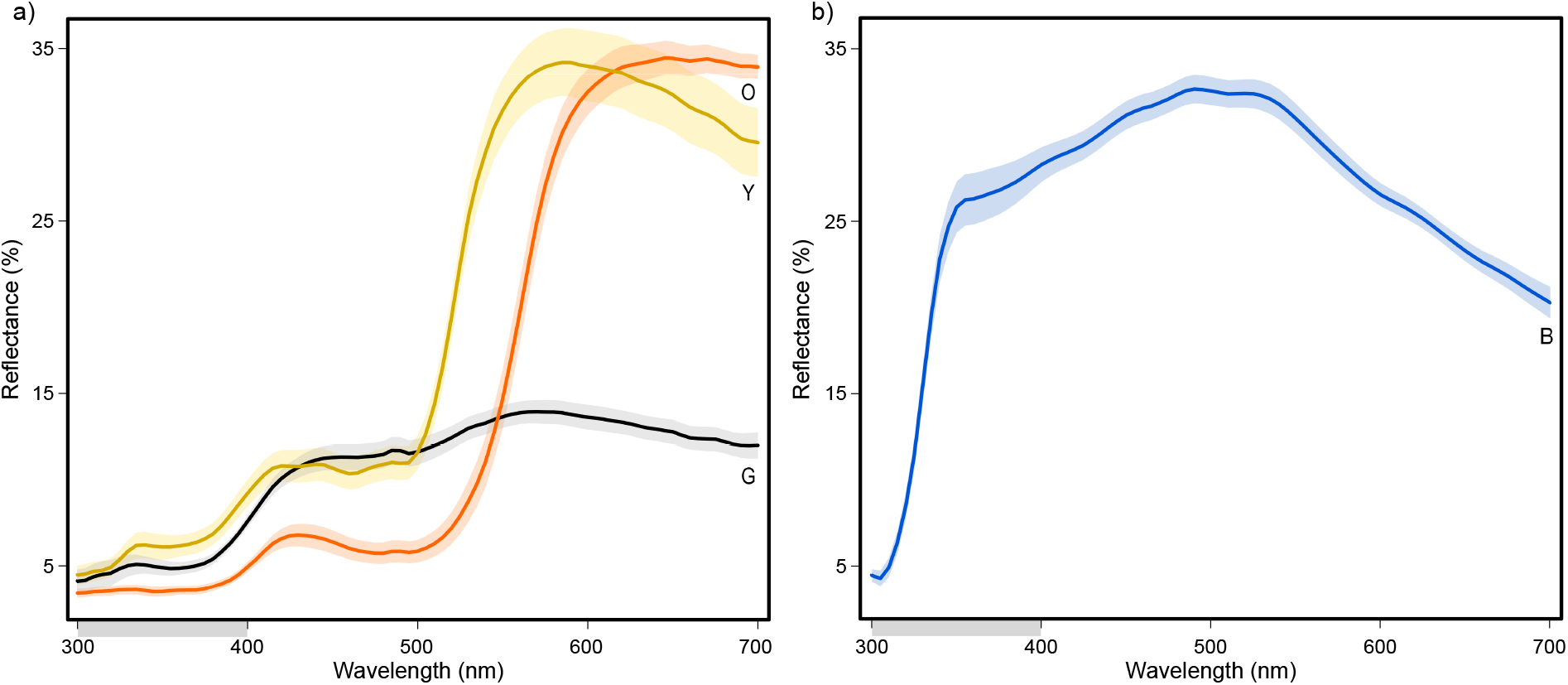
Average spectral reflectances of male throat colours found in (a) *C. modestus* measured from Southern Flinders Ranges populations: orange (O), yellow (Y), grey (G), and (b) *C. decresii* ultra-violet blue throat colour measured from Mainland South populations. Standard error shown as shaded region (N = 10 for each colour). Ultraviolet wavelengths are shown with a grey bar on the *x*-axis.

**Figure 2.**
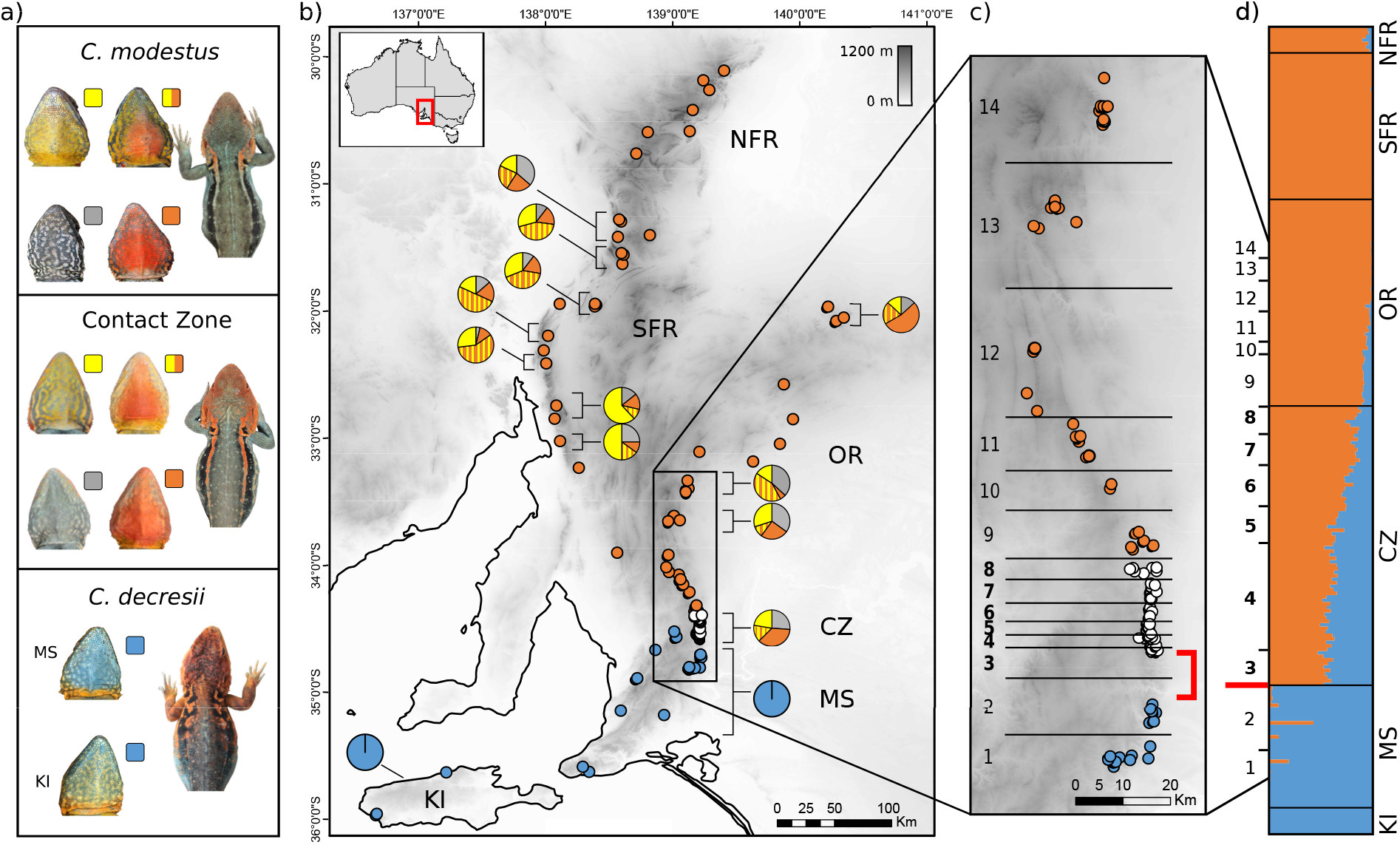
Geographic variation in throat colour morphs and genetic structure between *C. modestus* and *C. decresii*. (a) Representative throats (*C. modestus*: yellow, orange-yellow, grey and orange; *C. decresii*: blue) and dorsolateral phenotypes found in each species and the contact zone. (b) Map showing sampling localities of individuals used in genetic analyses: *C. modestus* (orange circles; Northern Flinders Ranges [NFR], Southern Flinders Ranges [SFR], Olary Ranges [OR]), *C. decresii* (blue circles; Mainland South [MS], and Kangaroo Island [KI]), and genetically admixed individuals (white circles; 0.1 < *q* < 0.9). Pie charts indicate relative frequencies of male throat colour morphs. Proportions for the Southern Flinders Ranges and northernmost population in the Olary Ranges are adapted from McLean, Stuart-Fox, & Moussalli, 2015. Throat colour morph data are not available for the Northern Flinders Ranges. Elevation in meters is represented by grey shading on the scale bar. (c) An enlarged map section showing sites across the contact zone transect used for geographic cline analyses; (d) Results of a Bayesian analysis of ancestry in the program STRUCTURE (K = 2). Each horizontal bar represents an individual and individuals are ordered by latitude and population. The proportion of orange and blue represents the proportion of *C. modestus* and *C. decresii* ancestry in each individual. A density trough between sites 2 and 3, where surveys did not recover any individuals, is indicated in red in panels B and C.

The throat colour polymorphism is likely to have evolved in *C. decresii* sensu lato and then was subsequently lost in *C. decresii* sensu stricto, thereby facilitating the evolution and fixation of the novel blue morph (McLean, Stuart-Fox, et al., 2014). Genetic divergence between the species (3.7% net sequence divergence in mtDNA) is consistent with range contraction to allopatric refugia during glacial-interglacial Pleistocene cycles (Byrne, 2008; McLean, Stuart-Fox, et al., 2014), where differing site-specific selective pressures may have led to the evolution of divergent phenotypes. A small number of genetically admixed individuals have been identified where the species meet geographically, suggesting that subsequent post-glacial range expansion has resulted in a secondary contact zone (McLean, Stuart-Fox, et al., 2014).

This is a compelling study system to explore the evolutionary consequences of hybridisation between colour polymorphic and monomorphic species which differ in morph number and type. Furthermore, the biology of both species and the polymorphism of *C. modestus* are well-characterised. *Ctenophorus modestus* and *C. decresii* are sexually dichromatic: females have cream coloured throats with a cryptic dorsal phenotype. Throat and dorsolateral coloration are important intraspecific sexual signals, which are emphasised during complex movement-based territorial and courtship displays (Gibbons, 1977, 1979; Osborne, Umbers, Backwell, & Keogh, 2012; Ramos & Peters, 2016; Stuart-Fox & Johnston, 2005). Male throat coloration develops at sexual maturity (ca. 18 months), is fixed for life, and autosomally inherited. The four morphs of *C. modestus* employ alternative behavioural strategies (Yewers, Pryke, & Stuart-Fox, 2016) corresponding to differing hormonal profiles (Yewers, Jessop, & Stuart-Fox, 2017), and are associated with MHC supertypes (Hacking, Stuart-Fox, Godfrey, & Gardner, 2018). The discrete polymorphism is likely controlled by two independently segregating loci, though the extent of orange or yellow within morphs behaves as a quantitative (polygenic) trait (Rankin, McLean, Kemp, & Stuart-Fox, 2016).

We used a method of double digest restriction-site associated DNA sequencing to identify single nucleotide polymorphisms (SNPs), sequenced the mitochondrial ND4 gene, and characterised male throat and dorsolateral phenotype across a naturally occurring contact zone between the polymorphic *C. modestus* and monomorphic *C. decresii*. Using geographic cline analysis, we compared the frequency of these genetic and phenotypic traits along a transect through the contact zone from pure *C. modestus* to pure *C. decresii* populations. Furthermore, we conducted captive-breeding to investigate F1 hybrid phenotype independent of exogenous selection. Specifically, we examined i) the geographic extent of hybridisation, ii) whether hybridisation is due to recent or earlier admixture, iii) whether hybridisation is bidirectional or asymmetric, and iv) whether there is evidence of selection on phenotype. We predicted that selection on throat coloration to maintain the *C. modestus* polymorphism should result in steeper and narrower clines compared to SNP and dorsolateral phenotype clines.

## Materials and Methods

### Field sampling

*Ctenophorus modestus* and *C. decresii* are small [snout-vent-length (SVL) ≤ 90 mm] diurnal rock-dwelling agamids endemic to South Australia. *Ctenophorus modestus* occurs in the Flinders and Olary Ranges whereas *C. decresii* occurs in the Mount Lofty Ranges, Fleurieu Peninsula, and on Kangaroo Island (Figure 2b). The contact zone is found in an area of relatively low-lying grassland between the rocky ranges of the parental species, coinciding with a change from relatively temperate to more semiarid conditions (Houston 1974; McLean et al. 2014b). Our sampling design aimed to capture the full geographic range of both species (Figure 2b, Table S1) to reaffirm previously described genetic divergence based on multi-loci sequence data and microsatellites (McLean, Stuart-Fox, et al., 2014). To extend this previous work we intensively sampled 14 sites along a north-south 160 km transect through the contact zone (Figure 2c, Table S2). As the geographic extent of hybridisation was initially unknown, the transect was anchored on either end by two genetically pure *C. decresii* (southernmost sites 1 – 2) and *C. modestus* (northernmost sites 13 – 14) populations, which were geographically connected along a rock range. Sites were demarcated by geographic distance when possible; however, areas with high densities of sampling were demarcated more finely to allow greater accuracy in analyses. Lizards were captured with a lasso (telescopic pole and fishing line) or by hand and released at the site of capture following data collection. A genetic sample (blood or tail tip stored in 99% ethanol at −20°C) and phenotypic data (described below and in Supplementary Information) were taken from each individual. We focused on collecting data from adult males (SVL ≥ 65 mm) and collected genetic samples from females opportunistically. Our dataset comprised 234 genetic samples in total from across both species’ full ranges (Table S1, Figure 2b). This included 152 genetic samples from the 14 sites along the contact zone transect, from which we also obtained phenotypic data from up to 233 males along the contact zone transect, depending on the trait (Table S2).

### SNP discovery

Genomic DNA was isolated and purified from blood or tissue using either an E.Z.N.A. Tissue DNA Kit (Omega Bio-tek, Norcross, GA, USA) or a GenCatch Blood and Tissue Genomic Mini-Prep Kit (Epoch Life Sciences, Sugar Land, TX, USA). DNA was sent to Diversity Arrays Technology (Bruce, ACT, Australia) for SNP genotyping using DArTSeq, a genome complexity reduction method similar to double digest restriction-site associated DNA (ddRAD) sequencing protocol. Protocols are described in detail for broad applications in Kilian et al., 2012 and specifically for the genus *Ctenophorus* in Melville et al., 2017. Briefly, genomic DNA was digested with two restriction endonucleases (PstI/HpaII), PCR amplified (1 min at 94°C; 30 cycles of 94°C for 20 s, 58°C for 30 s, 72°C for 45 s; final extension at 72°C for 7 min), and sequenced on an Illumina HiSeq2500. ‘High-density’ DArTseq SNP arrays use 2,500,000 (±7%) reads per sample in marker calling. Two technical replicates of each DNA sample were genotyped to assess reproducibility. DArT analysis pipelines and a calling algorithm (DArTsoft14) were used to process sequences into a matrix of SNP loci. As reference genomes, we used the *C. decresii* sensu lato draft genome (McLean, Stuart-Fox, and Moussalli, unpublished data) and the fully annotated central bearded dragon genome (*Pogona vitticeps;* Georges et al., 2015). In total, we obtained 172,893 SNPs from 234 individuals with 41.54% missing data. Using the R package *dartR* (v1.1.6, Gruber et al. 2018), we filtered this dataset conservatively with thresholds of 100% reproducibility, ≤1% missing data per locus, ≤10% missing data per individual, and also removed loci that deviated from Hardy-Weinberg Equilibrium (*P* < 0.05). This resulted in a dataset of 1,333 SNPs from 230 individuals with 0.26% missing data. Additionally, to evaluate potential bias associated with stringent filtering, we repeated the filtering steps with a threshold of ≤5% missing data per locus (other thresholds unchanged), resulting in a dataset of 6,889 SNPs from 225 individuals with 1.46% missing data. Analyses run on this dataset indicated qualitatively similar results (Figure S1).

### Mitochondrial DNA

We sequenced an 850 base-pair region of the mitochondrial genome which included the NADH dehydrogenase subunit four (ND4), and tRNAhis, tRNAser and tRNAleu regions using the primers ND4F (Forstner, Davis, & Arèvalo, 1995) and tRNA-Leu (Scott & Keogh, 2000) for a subset of 74 individuals from sites across the contact zone transect (Table S1). PCR was performed following procedures in McLean, Stuart-Fox, et al., 2014. PCR products were submitted to the Australian Genome Research Facility (Melbourne, Victoria, AUS) for purification and forward and reverse sequencing on an Applied Biosystems 3730xl DNA Analyzer. Sequences were edited and aligned with additional ND4 sequences spanning the range of *C. modestus* and *C. decresii* (McLean, Stuart-Fox, et al., 2014) in Geneious (v6.1.7; Kearse et al. 2012). Haplotype networks were constructed using the statistical parsimony algorithms of TCS (v1.21; Clement et al. 2000).

### Quantifying population structure and admixture

We examined genome-wide patterns of divergence using the Bayesian model-based clustering approach implemented in STRUCTURE (v2.3.4; Pritchard, Stephens, & Donnelly, 2000). After an initial burn-in of 500,000 Markov chain Monte Carlo (MCMC) generations, we ran 500,000 generations with 10 independent replicates for K values ranging from 1 to 5 under the “admixture” model with correlated allele frequencies and prior population information, all other parameters were kept at default, and convergence of runs was visually assessed. We used STRUCTURE HARVESTER (Earl & VonHoldt, 2012) to identify the optimum value of K for our dataset and summarised results across the selected value of K using CLUMPP (v1.1.2; Jakobsson & Rosenberg, 2007). We used the Bayesian estimation of the *q*-value (proportion of an individual’s genome inherited from each parental population) with 95% confidence intervals (CI) as a hybrid index. Individuals were identified as pure parentals or putative hybrids based on their hybrid index and 95% CI: an individual was classified as parentals if *q* was less than 0.1 (i.e. *C. modestus*) or greater than 0.9 (i.e. *C. decresii*), or of mixed ancestry if *q* was between 0.1 and 0.9.

We used previously identified population groups within each parental lineage to further examine population structure (*C. modestus*: Northern Flinders Ranges [NFR], Southern Flinders Ranges [SFR], Olary Ranges [OR]; *C. decresii*: Mainland South [MS], Kangaroo Island [KI]; McLean et al. 2014b) and Contact Zone (CZ). We calculated expected and observed heterozygosity, the number of alleles and allelic richness, estimated population pairwise F_ST_, and the inbreeding coefficient (F) for each population using the R packages *dartR* (v1.1.6; Gruber et al. 2018) and *adegenet* (v2.1.1; Jombart & Ahmed, 2011). We then performed a principal coordinate analysis (PCoA) using *dartR* to visualize pairwise genetic distances between individuals.

### Hybrid classification

We subsampled our SNP dataset to obtain a dataset of 200 loci with the highest average polymorphic information content values using the R package *dartR* (v1.1.6; Gruber et al. 2018). The dataset was analysed three times in parallel, each with a total of 100,000 MCMC generations after an initial burn-in of 50,000 generations, using the program NewHybrids (v1.1 Anderson & Thompson, 2002) and the R package *parallelnewhybrid* (Wringe, Stanley, Jeffery, Anderson, & Bradbury, 2017). We determined the probability that individuals within the contact zone belonged to one of the six genotypic classes resulting from the first two generations of hybridisation: parental forms (P1 and P2), F1 (P1 × P2), F2 (F1 × F1), and backcrossed (F1 × P1/P2). Although it is not always possible to unambiguously assign individuals into discrete classes due to the continuum that may be found in wild populations, classification may be suggestive of situations where contact is very recent or where F1 hybrids are infertile based on the absence of F2 hybrids and backcrosses, or where contact is several generations old based on the absence of pure parentals and F1 hybrids.

### Inferring population history

We estimated population demographics with the program δaδi v2.0.6 (Gutenkunst, Hernandez, Williamson, & Bustamante, 2009, 2010), which uses a diffusion approximation method to analyse joint site frequency spectra. For this analysis we considered only geographically connected populations of *C. modestus* and *C. decresii*, and therefore excluded NFR and KI from the dataset (Figure 2). The full set of SNPs (i.e. 172,893 SNPs from 234 individuals) was filtered with a threshold of ≤5% missing data per locus, ≤10% missing data per individual and 100% reproducibility, and secondaries were removed (i.e. retained one SNP per locus) to reduce linkage disequilibrium (filtering performed in *dartR* (v1.1.6, Gruber et al. 2018). This resulted in a dataset of 4,956 SNPs from 212 individuals with 1.48% missing data, which we converted to δaδi input format using the R package *radiator* (v1.1.5; Gosselin, Lamothe, Devloo-Delva, & Grewe, 2020).

We implemented a modified version of the model optimisation workflow (dadi_pipeline) from Portik et al., 2017. We were specifically interested in inferring the age of the contact zone and therefore tested two biologically realistic models of population history consistent with the observed structure of the contact zone: 1) “sec_contact_asym_mig”, where the two species split with no migration (T1), followed by a period of asymmetrical migration (T2; Figure 4); and 2) “sec_contact_asym_mig_three_epoch”, a more complex model involving a period of isolation after secondary contact (T3; Figure S2). Both models considered two geographically structured ‘populations’, on either side of the density trough between sites 2 and 3 (Figure 2): 1) *C. decresii* (MS; Figure 2) and 2) *C. modestus* pooled with admixed individuals (OR, SFR and CZ; Figure 2). We used folded site frequency spectra, a grid size of 200, 220, 240 and projection sizes of 60 and 180 alleles for *C. decresii* and *C. modestus* respectively. For each model, we performed two independent runs with four consecutive rounds of optimisation, with 100 replicates per round, increasing maximum iterations (maxiter = 5, 5, 10, 20), and decreasing fold in parameter generation (fold = 3, 2, 2, 1). Models were compared using the log-likelihood and Akaike information criterion (AIC), and the two independent runs converged on similar parameters (Table S5). We converted parameters from the top model as follows: reference effective population size as 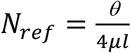, where *θ* was the scaled population parameter, μ was a mutation rate of 9 ×10^−10^ (Singhal & Moritz, 2013) and *l* was the length of analysed sequence (4956 SNPs × 70b = 34.7Mb); population size as *Nu* = *N*_*ref*_ × *nu*; time as *T* = *t*2*N*_*ref*_ × *g* where g was a generation time of 2 years; and migration (individuals per generation) as 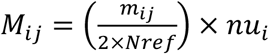.

### Captive-breeding and testosterone-induced expression of colour traits

To evaluate F1 hybrid phenotype independent of extrinsic selection pressures, we conducted captive breeding using 36 pure *C. decresii* individuals (female = 17, male = 19) collected from cline site 1 (southernmost site, longitude: 139.103°, latitude: −33.443°) and 41 pure *C. modestus* individuals (female = 20, male = 21) from cline site 14 (northernmost site, longitude: 139.103°, latitude: −33.443°; Table S2, Figure 2c) to produce reciprocal cross hybrids, in addition to pure offspring for comparison. Offspring were housed individually and reared under laboratory conditions (total = 138; southern = 11, northern = 54, hybrids = 73). We genotyped adults and offspring at five microsatellite loci (Am41, Cp10, Ctde45, Ctde05, Ctde12; Schwartz et al. 2007; McLean et al. 2014b) and confirmed paternity using a maximum likelihood approach in CERVUS (v3.0.7; Kalinowski, Taper, & Marshall, 2007; Marshall, Slate, Kruuk, & Pemberton, 1998). For all offspring, we induced throat colour expression by artificially elevating testosterone levels at nine months of age in both sexes (female = 64, male = 74; supplemental information). Females can be reliably induced to express discrete throat morphs found in males using testosterone because throat coloration is autosomally inherited; testosterone may influence the intensity of coloration but morphs remain objectively classifiable (Rankin & Stuart-Fox, 2015). Dorsolateral colour patterns remained unchanged in testosterone-treated females therefore dorsolateral phenotype was analysed only for male offspring. See Supplemental Information for full details of breeding, rearing, genotyping, and throat colour expression procedures.

### Analysis of phenotypic data

To characterise the phenotype of wild-caught males and captive-bred F1 hybrids, we took standardised ventral (i.e. throat) and dorsal photographs and spectral reflectance measurements of throat colours (details in Supplementary Information). The standardised photos were used to classify throats into existing morphs (orange, yellow, orange-yellow, grey, or blue), and to quantify the proportion of orange and yellow coloration present on the head and neck from dorsal photographs. We also manually quantified the number of distinct breaks in the dorsolateral stripe near the neck by visually inspecting all photographs of the dorsal region. Spectral reflectance measurements (300 – 700 nm) were used to estimate the receptor quantum catch (QC), the stimulation of retinal photoreceptors, for the violet to UV wavelength sensitive (UVS) photoreceptor because we were specifically interested in the diagnostic UV-blue component of throat coloration.

We performed principal component analyses (PCA) to explore variation in throat coloration variables (proportion of orange, proportion of yellow, proportion of blue) and dorsolateral phenotype variables (proportion orange, proportion of yellow, breaks in the dorsolateral stripe) in the R package *FactoMineR* (v1.4.1; Le, Josse, & Husson, 2008). In each, the lineages were differentiated along the first dimension (PC1) and this was used to collapse variables into a single measure of overall phenotype and retained for use in subsequent cline analyses. The quantum catch of the UV sensitive photoreceptor (QC_UVS_) was used singly in cline analyses to assess its frequency across the contact zone independent of throat colour morph. Due to missing data, 204 individuals were used in throat phenotype analyses whereas 233 individuals were used in dorsolateral phenotype analyses. Linear discriminant analyses were performed on pure individuals in the R package *MASS* (v7.3.50; Ripley, 2002) using all throat phenotype variables (proportion of orange, yellow, and blue, and QC_UVS_) and dorsolateral phenotype variables as independent variables.

### Geographic cline analysis

We used the R package *HZAR* (v0.2-5, Derryberry, Derryberry, Maley, & Brumfield, 2014) to fit a series of equilibrium geographic cline models (Gay, Crochet, Bell, & Lenormand, 2008; Szymura & Barton, 1986; Szymura & Barton, 1991) to both the molecular and phenotypic datasets, and in all cases using 1,000,000 MCMC generations following an initial burn-in of 100,000 generations. Specifically, we estimated cline models for the hybrid index (H_Index_; Bayesian *q*-value based on 1,333 loci), mtDNA haplotype frequencies, quantum catch of the UV wavelength sensitive receptor (QC_UVS_), PC1 for throat coloration (T_PC1_), and PC1 for dorsolateral phenotype (D_PC1_).

*HZAR* uses logistic regression equations to fit sigmoidal clines in a maximum-likelihood framework and describes cline centre (*c*) as the position where estimated trait frequency is 0.5 and width (*w*) as 1/slope at the cline centre. For molecular clines, we compared 15 models that varied in the fitting of trait interval (*p*; minimum and maximum frequencies at tails; fixed at 0 and 1, observed values, estimated values) and exponential decay tails (none, right only, left only, mirrored, both estimated separately). These 15 models were compared to an additional null model. For phenotypic trait data we compared models that varied in the fitting of exponential decay tails (none, right only, left only, mirrored, both estimated separately) and estimated mean and variance on the ends and in the centre. All models estimated cline centre and width as well as tail parameters if applicable (δ, distance from the centre to the tail; τ, tail slope). QC_UVS_ and T_PC1_ were normalised to values between 0 and 1 for plotting purposes.

To address model selection uncertainty, we conducted model averaging in an Akaike’s information criterion (AIC) framework. We calculated the Akaike weight (AIC*w*), a value between 0 and 1 which can be interpreted as the probability of being the best approximating model (Burnham & Anderson, 2002). Models were then ranked by AIC*w* downwards until the cumulative AIC*w* reached 0.95 to attain a 95% confidence set of best-ranked models (Lukacs, Burnham, & Anderson, 2010; Symonds & Moussalli, 2011). We recalculated the AIC*w* of models in the 95% confidence set and used those to calculate the model-averaged estimate of parameters (e.g. centre and width). We produced model-averaged clines by first multiplying the estimate for a given distance interval by the AIC*w*. This was done for each model within the 95% confident set of models. The AIC model averaged estimate was then obtained by summing up weighted estimates for each distance interval. Similarly, we multiplied the minimum and maximum estimate bounds by the AIC*w*, followed by summation to represent the credible region. Finally, the coincidence of cline centres was assessed by comparing model-averaged maximum-likelihood derived CIs (two log-likelihood unit support limits) of cline centres, with non-overlapping CIs considered statistically supported differences. Akaike weights were calculated using the R package *MuMIn* (v1.43.6; Bartoń 2018).

## Results

### Population structure and admixture

We found distinct population genetic structure within *C. modestus* and *C. decresii*, and evidence for the presence of hybrids in the contact zone. Based on the STRUCTURE analysis, only two clusters were identified, corresponding to the major split between *C. modestus* and *C. decresii* (Figure 2d, Table S3). The *q*-values ranged from 0.999 (pure southern) to 0.001 (pure northern). Across the contact zone transect, individuals from the southernmost sites 1 – 2 had a high probability of belonging to the *C. decresii* genetic cluster (*q* ≥ 0.90; mean ± SD = 0.99 ± 0.02) with the exception of two admixed individuals (*q* = 0.81 and 0.57). All individuals from the northernmost sites 9 – 14 had a high probability of belonging to the *C. modestus* genetic cluster (*q* ≤ 0.10; mean ± SD = 0.04 ± 0.03). Hybrids were identified within sites 3 – 8 where all individuals were of mixed ancestry and exhibited a highly skewed continuum of *q*-values ranging from 0.53 to 0.11. These sites will hereinafter be referred to as ‘the contact zone’, encompassing a distance of approximately 22.5 km. We also identified a density trough of approximately 12 km between sites 2 and 3 (Figures 2c–d) across which the genotype changes abruptly and multi-year surveys recovered no individuals. With the exception of two individuals at site 3, all admixed individuals in sites 3 – 13 had *C. modestus* mtDNA haplotypes. The *C. decresii* mtDNA haplotype was fixed in sites 1 – 2, including the only two admixed individuals identified in the geographic range of *C. decresii* (Figure S3).

The first axis of the PCoA constructed from 1,333 loci clustered all populations into two major groups corresponding to *C. modestus* and *C. decresii* (variance explained = 19.4%; Figure 3). We found evidence for the presence of hybrids along this axis, with hybrids clustered more closely to the Olary Ranges (OR) population of *C. modestus*. Population pairwise measures of F_ST_ between the contact zone and *C. modestus* OR/SFR/NFR populations (F_ST_ range: 0.058 – 0.216; Table S4) were lower relative to measures of F_ST_ between the contact zone and *C. decresii* MS/KI populations (F_ST_ range: 0.203 – 0.293). The contact zone had the lowest inbreeding coefficient (F = 0.095) and the highest level of heterozygosity (H_exp_ = 0.084, H_obs_ = 0.079) relative to parental populations, with very similar allelic number and richness to the OR population (Table 1). The second PCoA axis (variance explained = 3.5%; Figure 3) showed population structure within each species, with the Northern Flinders Ranges (NFR) and Kangaroo Island (KI) populations showing differentiation from other populations of their respective species, and from each other (F_ST_ = 0.571; Table S4). This can likely be attributed to the fact that each represents the northernmost and southernmost populations of their species, and KI is an insular population.

**Figure 3.**
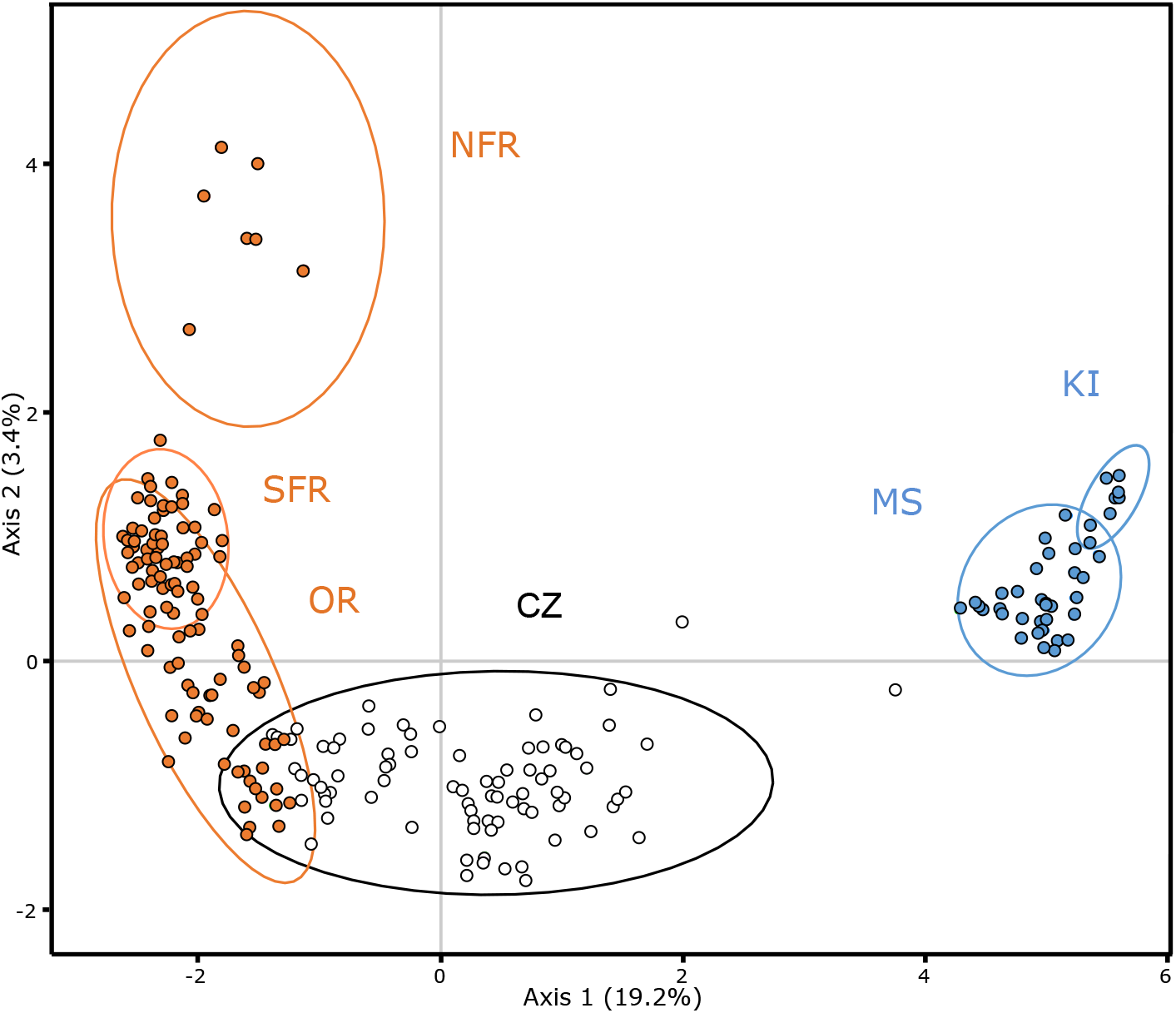
Two-dimensional principal coordinate plot (PCoA) constructed from the SNP data set (1,333 loci) showing pairwise genetic distances between individuals. 95% confidence ellipses of each group are shown (*C. modestus:* Northern Flinders Ranges [NFR], Southern Flinders Ranges [SFR], Olary Ranges [OR]; Contact Zone (CZ); *C. decresii* (Mainland South [MS], and Kangaroo Island [KI]). Two outlying admixed individuals (from the 95% confidence ellipse) were those found in the geographic range of *C. decresii* (MS).

**Table 1.**
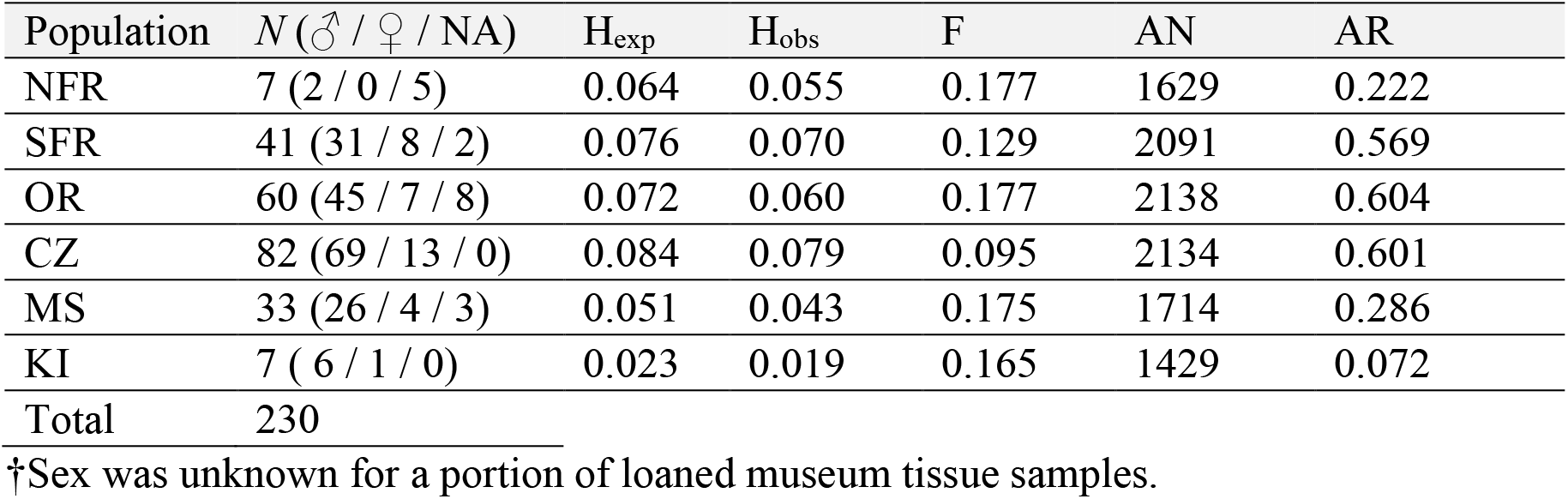
Populations sampled across the ranges of *C. modestus* (Northern Flinders Ranges [NFR), Southern Flinders Ranges [SFR], Olary Ranges [OR]), *C. decresii* (Mainland South [MS], and Kangaroo Island [KI]), and Contact Zone (CZ), with details of sample size for the SNP dataset (*N*), expected and observed heterozygosity (H_exp_, H_obs_), inbreeding coefficient (F), allele number (AN), and allelic richness (AR).

| Population | $N$ (♂ / ♀ / NA) | $H_{\text{exp}}$ | $H_{\text{obs}}$ | $F$ | AN | AR |
| --- | --- | --- | --- | --- | --- | --- |
| NFR | 7 (2 / 0 / 5) | 0.064 | 0.055 | 0.177 | 1629 | 0.222 |
| SFR | 41 (31 / 8 / 2) | 0.076 | 0.070 | 0.129 | 2091 | 0.569 |
| OR | 60 (45 / 7 / 8) | 0.072 | 0.060 | 0.177 | 2138 | 0.604 |
| CZ | 82 (69 / 13 / 0) | 0.084 | 0.079 | 0.095 | 2134 | 0.601 |
| MS | 33 (26 / 4 / 3) | 0.051 | 0.043 | 0.175 | 1714 | 0.286 |
| KI | 7 (6 / 1 / 0) | 0.023 | 0.019 | 0.165 | 1429 | 0.072 |
| Total | 230 |  |  |  |  |  |

†Sex was unknown for a portion of loaned museum tissue samples.

### Hybrid classification

We found a strong bias in genetic introgression with admixed individuals being predominantly backcrossed with *C. modestus*. Of individuals within the contact zone, 35% had intermediate *q*-values (0.4 < *q* < 0.6) and a large proportion (65%) had *q*-values between 0.39 and 0.10 suggestive of backcrossing. NewHybrids analysis classified all individuals of *C. modestus* and *C. decresii* (0.1 > *q* > 0.9) as pure parental forms (PP ≥ 99.1%). Further, 59 of 79 individuals in the contact zone were classified to one of the six classes with a posterior probability (PP) of ≥ 95%. Additionally, 6 individuals were classified as pure *C. modestus* (P1), 36 as F2 hybrids, and 19 as backcrosses to the northern lineage (B1). We did not recover any F1 hybrids or backcrosses to *C. decresii*, posterior probabilities for these two categories were uniformly low (average PP = 0.0%). The two admixed individuals found in the geographic range of *C. decresii* were ambiguously classified as an F2 (PP = 54.2%) and a backcross to *C. decresii* (PP = 80.8%). The 22 individuals unable to be confidently classified (PP ≤ 95%) likely represent the continuum present in wild populations, unsuited for discrete classification, and difficult to distinguish even with many diagnostic markers (Boecklen & Howard, 1997; Vähä & Primmer, 2006).

### Population history

The δaδi analysis showed that the model of secondary contact with ongoing asymmetrical migration (“sec_contact_asym_mig”) was more likely than the model with subsequent isolation (“sec_contact_asym_mig_three_epoch”), based on log-likelihood and AIC (Figure S4, Table S5). To aid interpretation, we converted parameter estimates into biologically meaningful values (Figure 4; Figure S2). For the top model, the total time since divergence of *C. decresii* and *C. modestus* was ~1.95 million years (T1+T2), with secondary contact occurring ~170,000 years ago (T2). The ‘population’ size of *C. modestus* (Nu2) was approximately 4 times larger than *C. decresii* (Nu1), and migration was asymmetrical with 0.36 individuals from *C. modestus* to *C. decresii* per generation compared to 1.55 individuals from *C. decresii* to *C. modestus* per generation (Figure 4).

**Figure 4.**
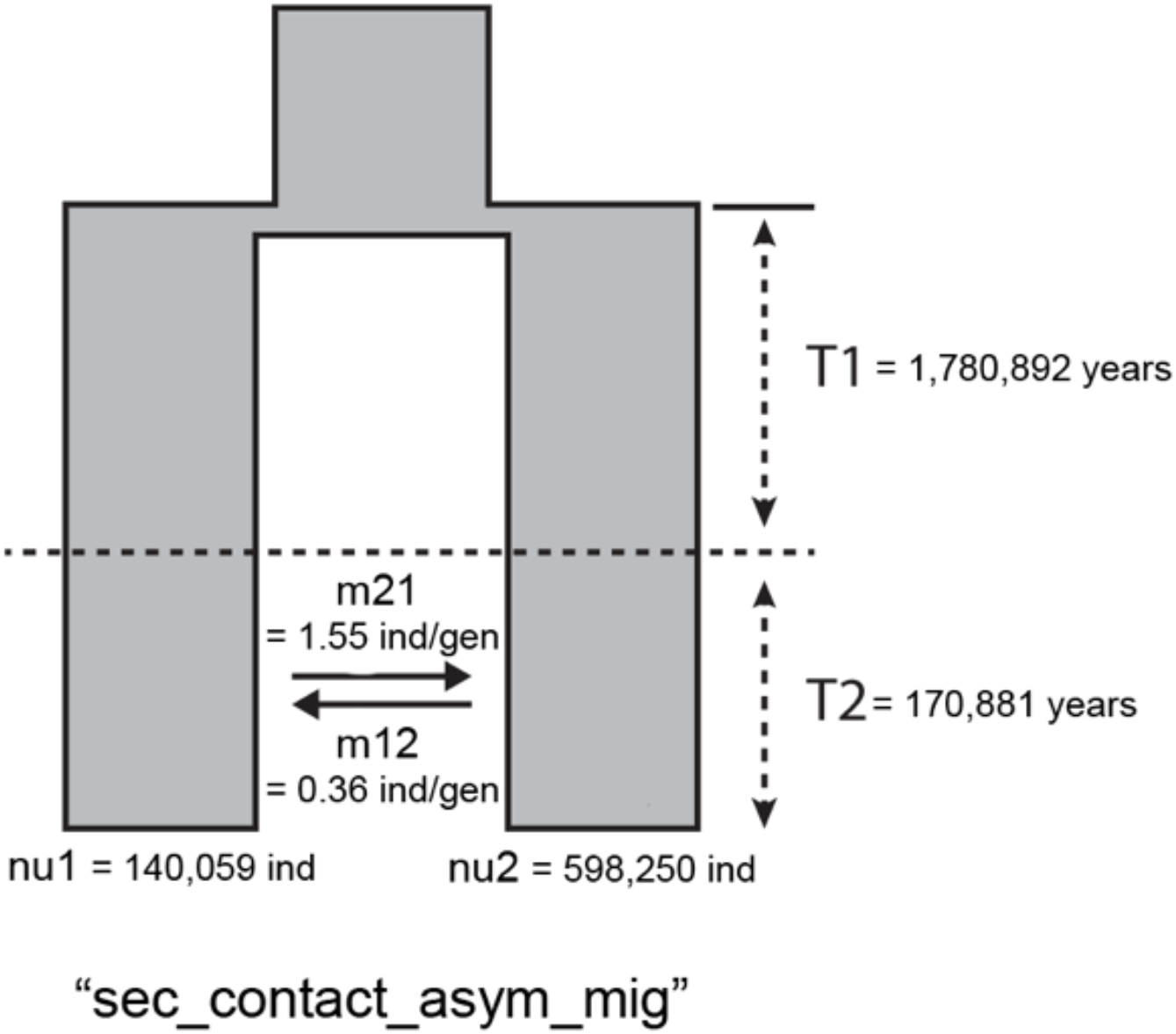
Schematic representation of the top demographic model from δaδi analysis. Populations 1 and 2 are *C. decresii* and *C. modestus* (pooled with admixed individuals) respectively. Converted values are provided for times (T1, T2, T3; in years), migration rates (m12, m21; in individuals per generation) and population sizes (nu1 and nu2; in number of individuals).

### Hybrid throat and dorsolateral phenotypes

The throat phenotype of wild-caught hybrid males was more similar to that of *C. modestus*, exhibiting all four throat colour morphs with uniformly low levels of UV reflectance. Linear discriminant analyses (LDA) on throat variables showed *C. modestus* and *C. decresii* strongly differentiated along LD1 which was driven primarily by the proportion of blue and secondarily by QC_UVS_ and the proportion of orange (variance explained = 99.8%; Figure 5a, Table S6). The 95% confidence ellipses drawn for the throat phenotypes of *C. modestus* and hybrid males overlie each other. The large majority of hybrid throats cluster with *C. modestus* throats. Blue coloration was not detected on the throats of pure *C. modestus* males. A degree of blue coloration (≥ 5% blue pixels in segmentation analysis), however, was found on a portion of hybrid throats of each morph: grey (47.4%, N = 19), orange (5.2%, N = 39), orange-yellow (6.7%, N = 15), yellow (11.8%, N = 17; Figure 6a). The throat phenotype of the two hybrid individuals found in the range of *C. decresii* had larger proportions of blue coloration on the throat relative to other hybrids.

**Figure 5.**
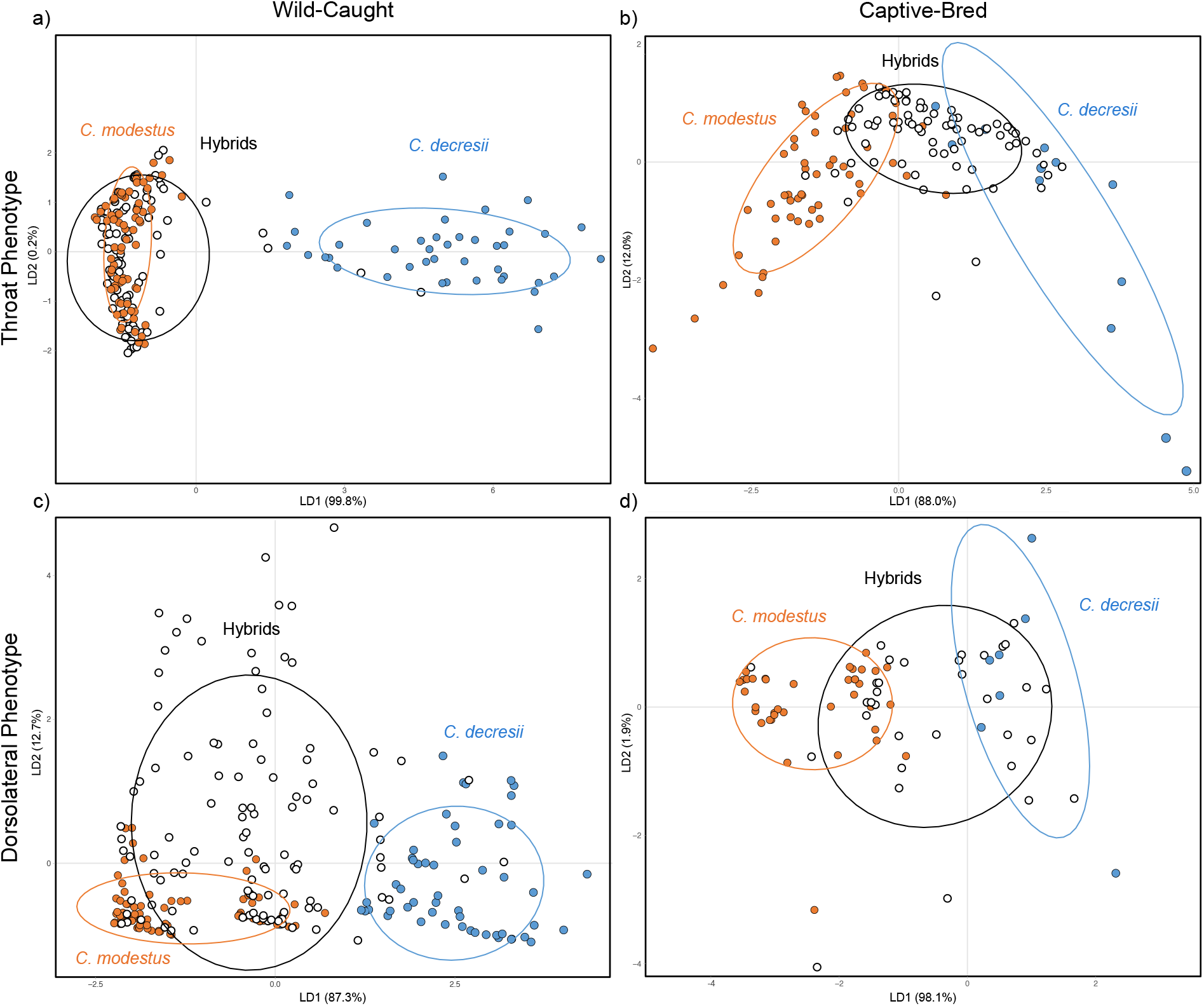
Biplots of linear discriminant analyses on (a, b) throat and (c, d) dorsolateral phenotype variables for (a, c) wild-caught individuals along the contact zone transect and (b, d) captive-bred pure and F1 hybrid offspring. 95% confidence ellipses are shown.

**Figure 6.**
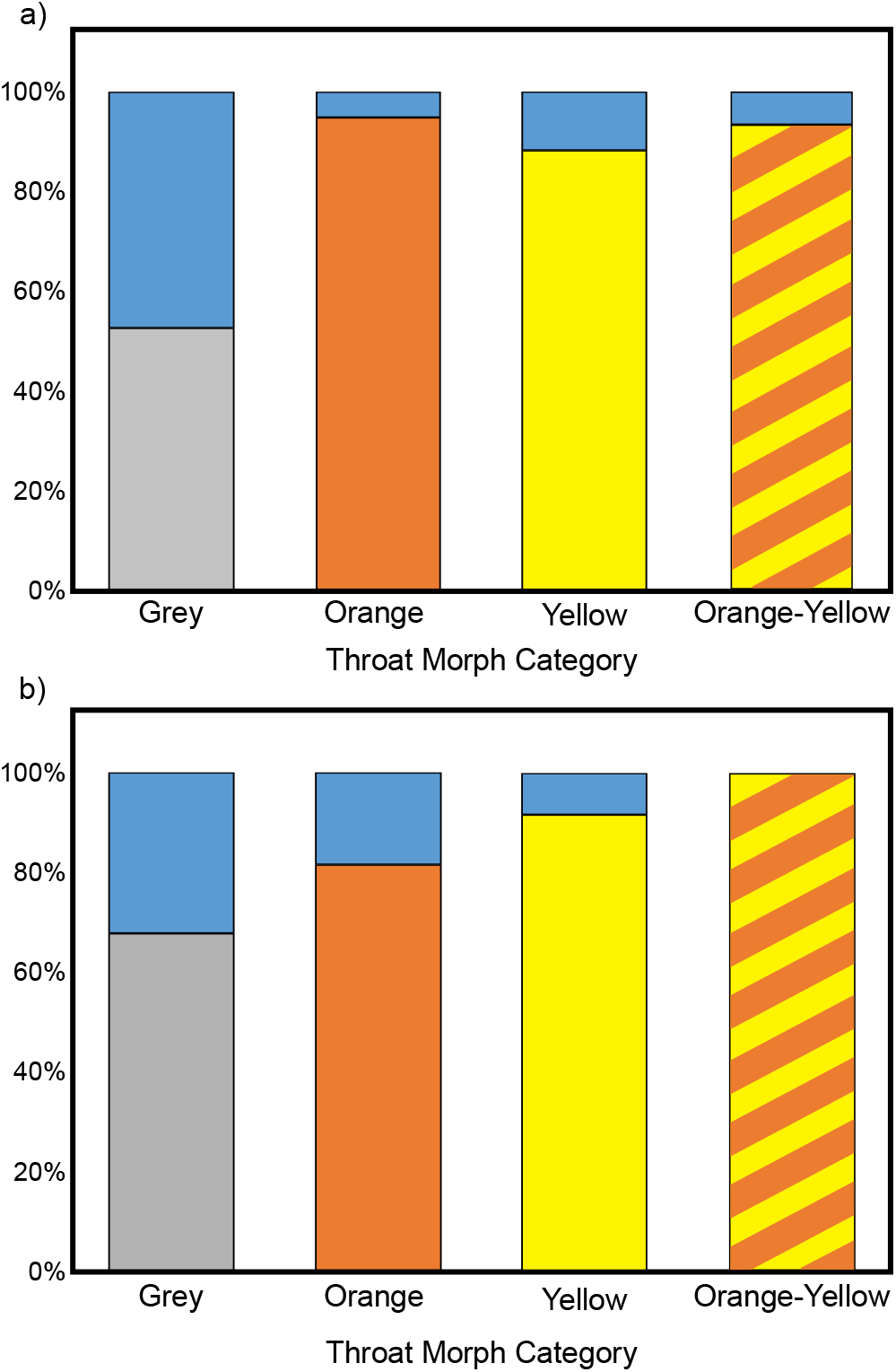
Percentage of individuals in each throat morph category which exhibit a degree of blue coloration on the throat for (a) wild-caught male hybrids (grey N = 19, orange N = 39, orange-yellow N = 15, yellow N = 17) and (b) captive-bred F1 hybrids (both sexes; grey N = 31, orange N = 27, orange-yellow N = 2, yellow N = 12).

In contrast to wild-caught hybrid throats, captive-bred F1 hybrid throat phenotype was more intermediate and exhibited a wide range of UV reflectance levels varying from those characteristic of *C. modestus* to *C. decresii*. Linear discriminant analysis on throat variables of captive-bred F1 hybrids showed strong differentiation between lineages along LD1 driven primarily by QC_UVS_ and secondarily by the proportions of blue and yellow (variance explained = 88.0%; Figure 5b, Table S6). The 95% confidence ellipse drawn for hybrid throats overlaps those of both *C. modestus* and *C. decresii* throats to a similar extent. Results were essentially unchanged when considering hybrids sired by a *C. modestus* or *C. decresii* male and by offspring sex. Similar to wild hybrids, all four *C. modestus* throat colour morphs were recovered in captive-bred hybrids, and all but the orange-yellow morph had a proportion of individuals with a blue component: grey (32.3%, N = 31), orange (18.5%, N = 27), orange-yellow (0%, N = 2), yellow (8.3%, N = 12; Figure 6b). Although the artificial elevation of testosterone levels has been shown to induce the expression of *C. modestus* throat colour morphs in juveniles and females (Rankin et al., 2016; Rankin & Stuart-Fox, 2015), we found that it did not comparably induce the expression of the blue throat colour morph in all pure *C. decresii* offspring. Therefore, the presence of blue coloration on F1 hybrid offspring throats may have been underestimated.

**Figure 7.**
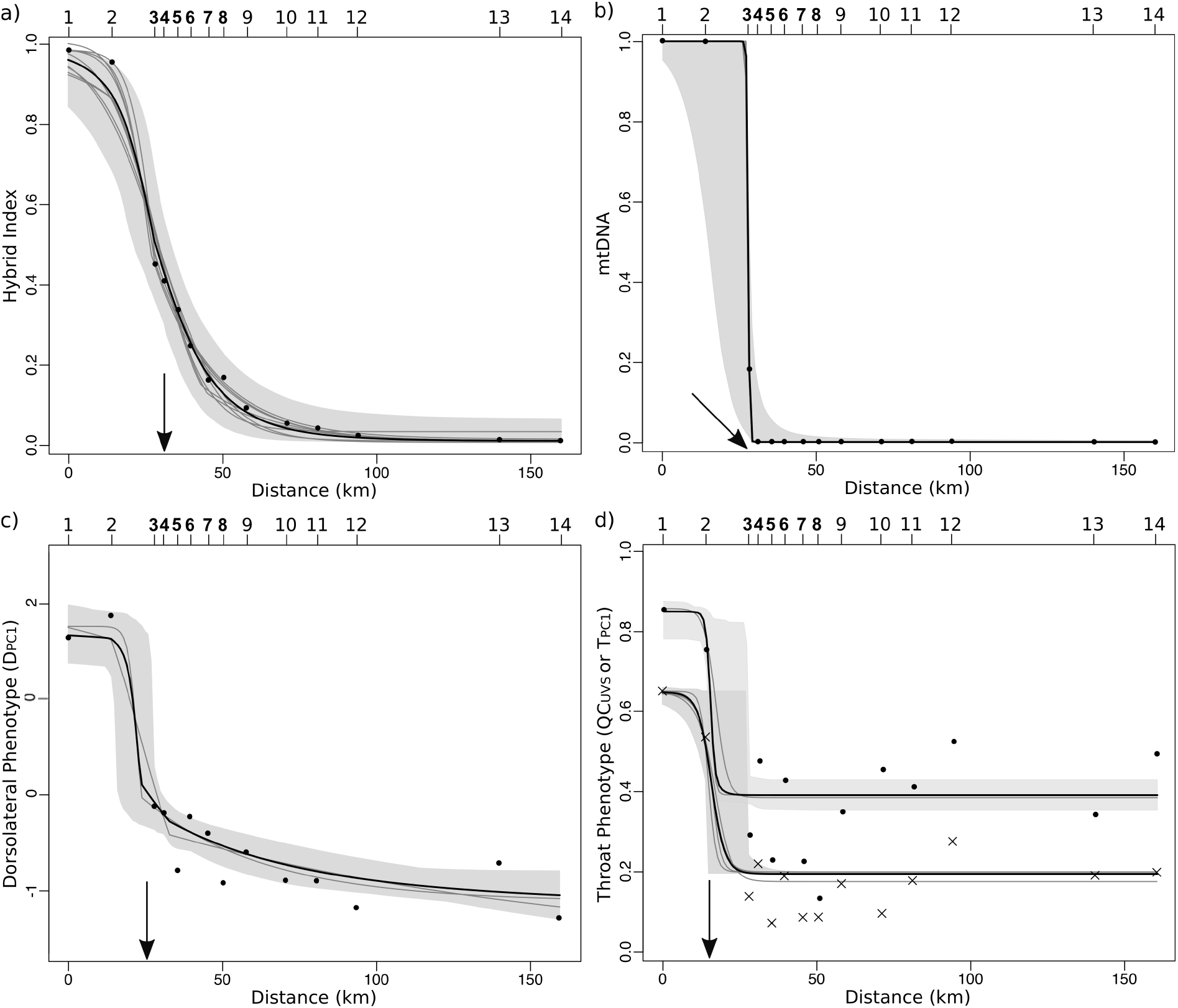
AICc model-averaged geographic clines (black) and credible regions, clines comprising the 95% confidence set of models (grey), and observed frequencies at each site for(a) the hybrid index estimated from 1333 loci (N = 154), (b) mtDNA haplotype frequencies (N = 74), (c) dorsolateral phenotype (D_PC1_; N = 233), and (d) throat phenotype clines (N = 204 for both): quantum catch of UV receptor (QC_UVS_; lower cline, frequencies shown with ‘×’ symbols) and throat colouration (T_PC1_; upper cline). Numbers along the top refer to sampling sites (see Table S2 for sample sizes at each site), bolded sites (3 – 8) comprise the contact zone. Transect distance is the cumulative distance from the southern-most population with increasing distance northwards. Arrows indicate cline centres; throat phenotype clines have essentially the same centres and are indicated with one arrow.

Dorsolateral phenotype in both wild-caught and captive-bred hybrids is intermediate between the lineages (Figure 5c–d). Within the contact zone, differentiation between lineages along LD1 is driven primarily by the number of breaks in the dorsolateral stripe and proportion of orange (Table S6). The 95% confidence ellipses drawn for the dorsolateral phenotypes of hybrids and *C. modestus* males overlap incompletely, likely reflecting the high frequencies of backcrossing to *C. modestus*. In captive-bred hybrids, comparable results were found with differentiation driven by breaks in the lateral stripe (Table S6) and the 95% confidence ellipse for hybrid dorsolateral phenotype overlaps those of both *C. modestus* and *C. decresii* dorsolateral phenotypes.

### Geographic clines

Model-averaged clines for mtDNA and throat phenotype (QC_UVS_ and T_PC1_) exhibited relatively narrower clines with abrupt transitions suggesting selection on these traits, compared to clines for the hybrid index (H_Index_) and dorsolateral phenotype (D_PC1_; Table 2). The best-fitting clines for the hybrid index and mtDNA had coincident centres with overlapping CIs but showed discordance in slope where the mtDNA cline was significantly narrower (H_Index_: *c* = 28.41 km, *w* = 29.77 km; mtDNA: *c* = 27.78 km, *w* = 0.64 km; Figure 8a–b). The cline fitted for the dorsolateral phenotype (*c* = 23.02 km, *w* = 13.97) was coincident with that of the hybrid index (Figure 8c). Clines for throat phenotype (QC_UVS_ and T_PC1_) were similar in centre and width to each other (QC_UVS_: *c* = 15.90 km, *w* = 8.48 km; T_PC1_: *c* = 15.38 km, *w* = 3.22 km; Figure 8d), their centres were displaced by 12.51 and 13.03 km to the south from the hybrid index cline centre and do not have overlapping CIs with the hybrid index cline. See Table S7 for full details of cline models in the AIC 95% confidence set for each trait.

**Table 2.**
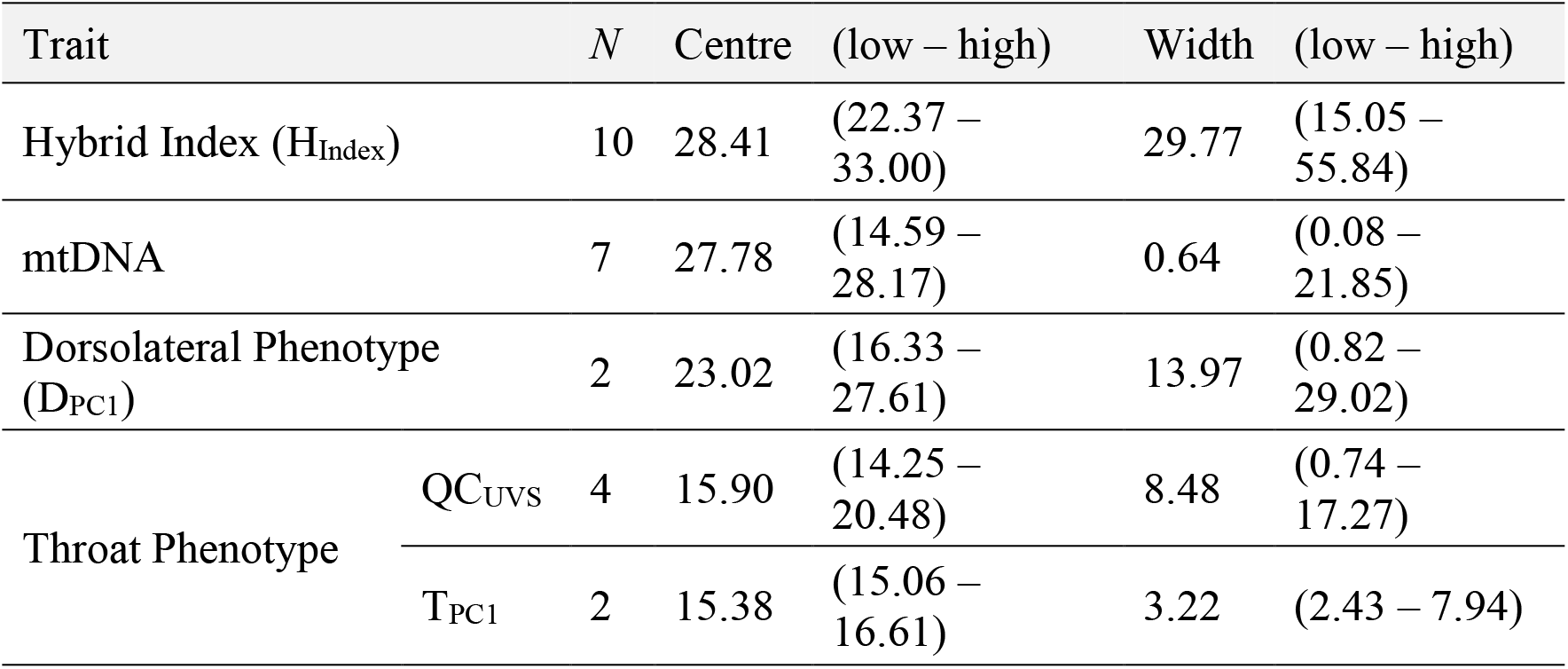
Model-averaged estimates for cline centre and width based on the AIC 95% confidence set models, with *N* representing the number of models within the AIC confident set. Cline centre is the distance (km) from the southern-most sampling site and width (*w*) is 1/slope at the centre. Two log-likelihood unit support limits are shown as ‘low’ and ‘high’. See Table S7 for full details of cline models.

## Discussion

Evaluating the evolutionary outcomes of secondary contact between lineages that differ in morph composition can provide valuable insights into the role of polymorphism in speciation. We characterised genomic and phenotypic patterns across the contact zone between the colour polymorphic *C. modestus* and monomorphic *C. decresii* and concurrently measured the phenotype of captive-bred F1 hybrids. We found that the contact zone is geographically narrow relative to the parental ranges (approx. 30 km) with no parental forms or F1 hybrids present in the genetic centre, suggesting that hybridisation is neither recent nor ongoing. Genomic introgression was asymmetric, with hybrids backcrossing almost exclusively to *C. modestus*. Formal comparison of two plausible models of population history for this system supports a prolonged period of secondary contact with asymmetrical migration. This suggests that despite potential for admixture throughout the last interglacial period, hybridisation between the species is not ongoing. Dorsolateral phenotype across the contact zone was consistent with patterns of genetic admixture, whereas the cline for throat phenotype was displaced relative to clines for the hybrid index and dorsolateral phenotype. Throat phenotype of wild-caught hybrids resembled that of *C. modestus*, with all four *C. modestus* throat morphs present and uniformly low UV reflectance. By contrast, both dorsolateral and throat phenotype were intermediate in captive-bred F1 hybrids. Taken together, these data are consistent with selection on throat phenotype across the contact zone.

The pattern of asymmetric genomic introgression, with limited backcrossing to the southern lineage, could reflect multiple evolutionary and demographic scenarios. First, extrinsic barriers may hinder dispersal and migration, in turn preventing backcrossing. Second, the pattern of asymmetric genomic introgression coupled with the prevalence of *C. modestus* mtDNA could potentially be explained by demographic circumstances. For instance, a greater abundance of *C. modestus* individuals during initial periods of hybridisation with continued asymmetric migration into the contact zone, or the sex-biased dispersal of *C. decresii* males to the north, could both result in asymmetric introgression. Third, intrinsic genomic incompatibilities could limit backcrossing via reinforcement (i.e. selection against maladaptive hybridsation). We discuss each explanation below in light of our data and available evidence.

The restricted gene flow to *C. decresii* is associated with an approximately 12 km density trough (Figure 3b–c). In this region, it appears that hybrid and/or pure populations either occur in levels below human detection or do not currently exist despite continuous rocky habitat with no obvious barriers to dispersal (e.g. major roads, waterways, human structures; Figure S5). The density trough was surveyed thoroughly in 2016 and 2017 and briefly in 2013. The presence of suitable habitat was confirmed by surveys on foot, satellite imagery (Figure S5), and/or flying drone surveys; this area is unlikely to constitute an ecological barrier. Furthermore, a model with prolonged secondary contact and ongoing asymmetric migration better fits the data than one with subsequent isolation. Secondary contact over at least the last interglacial period suggests a sufficient period for dispersal across the 12 km density trough in the absence of extrinsic barriers. Although dispersal has not been examined in *C. decresii*, dispersal estimates in other lizard species range from 0.01 km/gen to 0.1 km/gen for Dactyloidae and Lacertidae (Calsbeek, Duryea, Parker, & Cox, 2014; reviewed in McEntee et al., 2020; Vercken, Sinervo, & Clobert, 2012). With a generation time of two years, dispersal across this distance is consistent with the estimated age of the contact zone. In general, the width of the genetic contact zone (approx. 30 km), even taking into consideration the 12 km density trough, is relatively narrow relative to the parental ranges (Figure 2b). This distance is in accordance with cline widths estimated from genetic data in other lizards: Teiidae (3.2 – 93.9 km; Dessauer, Cole, & Townsend, 2000; Gifford, 2008), Dactyloidae (3 – 5 km; Case & Williams, 1984; Johansson, Surget-Groba, & Thorpe, 2008), Lacertidae (2.7 – 29.9 km; Miraldo, Faria, Hewitt, Paulo, & Emerson, 2013), Phrynosomatidae (0.8 – 30.9 km; Leaché & Cole, 2007; Leaché, Grummer, Harris, & Breckheimer, 2017; J. C. Marshall & Sites Jr., 2001; Sites, Barton, & Reed, 1995).

Demographic factors alone are similarly unlikely to explain the observed asymmetric introgression. The greater abundance and migration of *C. modestus* individuals into the contact zone should result in a greater proportion of *C. modestus* genomic background (*C. decresii* genomic signature quickly eroded by *C. modestus* contribution). However, we found close to equal genomic proportions in many hybrids in the genetic centre. Despite opportunity for ongoing migration, albeit asymmetrical, hybrids backcrossed to *C. decresii* were almost entirely absent. Male-biased dispersal could underlie population genetic structure consistent with our results (Jockusch & Wake, 2002; Petit & Excoffier, 2009); however, it is unlikely that male-biased dispersal would exclusively be observed in *C. decresii* and not in *C. modestus* given their polygynous mating systems (reviewed in Li & Kokko, 2019).

The absence of extrinsic barriers coupled with the narrower and more abrupt transition exhibited by the mtDNA cline relative to the hybrid index, and the rarity of admixed individuals containing the *C. decresii* mtDNA haplotype, point to a role for intrinsic genomic incompatibilities (e.g. Helbig, Salomon, Bensch, & Seibold, 2001; Miller, Lipshutz, Smith, & Bermingham, 2014). Captive breeding experiments between *C. modestus* and *C. decresii* show no significant prezygotic barriers to the formation of F1 hybrids with either parental mtDNA haplotype (Dong, Rankin, McLean, & Stuart-Fox, unpublished data), although their reproductive viability is unknown. However, stronger incompatibilities and hybrid breakdown are more frequently seen in F2 and later generation hybrids during the early stages of divergence. This has been described extensively for both Dobzhansky-Muller incompatibilities (reviewed in Coyne & Orr, 2004) and for cytonuclear incompatibilities (Ellison & Burton, 2008; Fishman & Willis, 2006; Meiklejohn et al., 2013; Niehusis, Judson, & Gadau, 2008). Overall, our results support the hypothesis that intrinsic factors contribute to limiting gene flow, and backcrossing to *C. decresii*, regardless of mtDNA haplotype, may result in offspring that are inviable, infertile, or suffer reduced fitness.

Phenotypic patterns recovered from within the contact zone could theoretically result from selection on the throat colour signal or neutral processes. We found that both throat phenotype clines were displaced relative to the genetic cline, in contrast to the dorsolateral phenotype cline. The displaced throat phenotype clines and prevalence of the *C. modestus* throat phenotype in the contact zone could theoretically arise from genetic dominance. In this scenario, captive-bred F1 hybrids would be expected to have largely *C. modestus* throat phenotypes which contrasts with the observed intermediate phenotypes. Another possibility is that this phenotypic pattern is due to extensive backcrossing to *C. modestus*. While some wild hybrids may have northern throat phenotypes because they are heavily backcrossed to *C. modestus*, the prevalence of *C. modestus* phenotype in hybrids with close to equal proportions of *C. modestus* and *C. decresii* genomic backgrounds points towards selection on throat colour.

We did not detect the loss of any *C. modestus* throat morphs across the contact zone or in F1 hybrids, indicating that co-adapted trait complexes of the *C. modestus* polymorphism are stable despite genetic disruption. The maintenance of all four morphs in all populations of *C. modestus* despite environmental selection on morph frequencies suggests that negative frequency dependent selection is a strong evolutionary force counteracting loss of polymorphism due to drift (McLean, Stuart-Fox, & Moussalli, 2015). Although hybrid throats more closely resembled *C. modestus* throat morphs, we observed a proportion of each morph which exhibited a degree of intermediate nature where an area of blue coloration was present. There was a notably higher proportion of grey morphs which had blue coloration present. This may be due to the similar mechanism by which grey and blue skin are produced in *C. modestus* and *C. decresii* (McLean, Lutz, Rankin, Stuart-Fox, & Moussalli, 2017). Nevertheless, we did not detect any hybrid throats that exhibited a true *C. decresii* throat phenotype with both predominantly blue throat coloration and a high level of UV reflectance. This, in addition to the variable presence of UV reflectance in captive-bred F1 hybrids, points toward the uncoupling of blue coloration and UV reflectance in hybrids. Hybridisation can generate novel phenotypes (e.g. blue without UV); for example, admixture between treefrog lineages which have red and red/blue legs result in purple legs which are produced from the directional introgression of colour loci inherited from parental lineages (Akopyan et al., 2020).

Selection on throat coloration could be driven by mating behaviour which evolved via reinforcement and/or microhabitat characteristics. Mate preference trials with captive *C. modestus* and *C. decresii* predict a greater frequency of mating between *C. decresii* females with *C. modestus* males than the reciprocal combination(McLean, Bartle, Dong, Rankin, & Stuart-Fox, 2020). This contrasts with patterns observed within the contact zone, indicating that prezygotic barriers between species have not evolved in allopatry. However, postzygotic barriers (inferred from low frequency of hybrids backcrossed to the south) should select for the formation of prezygotic barriers to prevent maladaptive hybridisation (i.e. reinforcement). Extrinsic factors could also contribute to the *C. modestus* throat phenotype having a competitive advantage in the contact zone, for example, due to a relatively higher conspicuousness to conspecifics. The throat coloration of each species is more conspicuous against the predominant background colours of native lichen in their respective ranges (McLean, Moussalli, & Stuart-Fox, 2014). Within the contact zone, lichen coloration is similar to that found in the range of *C. modestus*, predominantly pale green. Therefore, sexual selection may operate through male-male interactions or female choice to result in a reproductive advantage of the *C. modestus* throat phenotype as a more conspicuous and detectable visual signal (reviewed in Servedio & Boughman, 2017). Similarly, the greater overlap of hybrid phenotype (e.g. ventral coloration) with one parental species was associated with environmental variation in *Hyperolius* frogs (Bell & Irian, 2019). Selection has been shown to drive the rapid spread of an advantageous trait/phenotype across contact zones, and in some cases into the other parental lineage (Baldassarre, White, Karubian, & Webster, 2014; Pardo-Diaz et al., 2012; Stein & Uy, 2006; While et al., 2015). In contrast, the dorsolateral phenotype cline indicates that it is not under similarly strong selection and appears to play a lesser role in reproductive isolation in sympatry. Male dorsolateral coloration may be constrained as it is under both sexual and natural selection because of its dual function as a intraspecific signal and in minimising conspicuousness to predators against native lichen (McLean, Moussalli, et al., 2014; Ramos & Peters, 2016).

### Conclusions

In summary, our study suggests that sexual signal divergence and intrinsic genetic incompatibilities play an important role in restricting gene flow between the polymorphic *C. modestus* and monomorphic *C. decresii*. The species remain distinct despite continued migration and opportunity for gene flow. The stability of the *C. modestus* polymorphism despite genetic disruption is consistent with the theoretical genetic architecture of colour polymorphism and correlated traits, namely control by pleiotropic or tightly linked genes. Taken together, these results lend support to the view that polymorphism can promote speciation.

## Acknowledgements

This study was funded by the Australian Research Council (DP150101048 and DP1092908) to D.S-F., Holsworth Wildlife Endowment and The David Lachlan Hay Memorial Fund to C.M.D., and Holsworth Wildlife Endowment, Alfred Nicholas Fellowship and Nature Foundation South Australia to C.A.M. We thank Madeleine Yewers, Steven Mesis, Katrina Rankin and Anne Auslebrook for assistance; private landowners for kindly allowing land access; and South Australian Museum for providing tissue samples. Data collection was conducted with permits from the Department of Environment, Water and Natural Resources, South Australia (permit nos. E25861-1 and Q26428-3), the Department of Environment, Land, Water and Planning, Victoria (permit nos. 10007000 and 10007751) and with approval by The University of Melbourne Animals Ethics Committee (approval nos. 1011760.1 and 1413220.4), the Wildlife Ethics Committee of South Australia (approval nos. 18/2010 and 25/2015). We thank Ben Phillips and three anonymous reviewers for insightful comments, which helped to improve the manuscript.

## Data Accessibility

The DArTseq genotype data, mtDNA alignment, R code and morphological data that support the findings of this study are openly available in Dryad at http://doi.org[doi], reference number [number]. All mtDNA sequences are deposited in GenBank, accession numbers MW261595 - MW261739.

*Data will be made available on Dryad upon acceptance of the manuscript.*

## Author Contributions

All authors contributed to study design and field sampling. C.M.D. and C.A.M. performed laboratory work. C.M.D. conducted data analyses and drafted the manuscript. C.A.M, A.M. and D.S-F provided input on analyses, interpretation, and the manuscript. All authors read and approved the final manuscript.

