## Supplemental Information for "When polymorphism and monomorphism meet: discordant genomic and phenotypic clines across a lizard contact zone"

**Table of Contents**


| | |
|---|---|
| Table S1. Details of all individuals used in the study. | 2 – 9 |
| Table S2. The sampling transect across the contact zone. | 10 |
| Table S3. Summary of STRUCTURE results for the dataset of 1333 loci. | 11 |
| Table S4. Pairwise $F_{ST}$ values between populations. | 12 |
| Table S5. Likelihoods and parameters from δaδi. | 13 |
| Table S6. Results of throat and dorsolateral phenotype analyses. | 13 |
| Table S7. Parameter estimates for cline models in 95% confidence sets. | 15 – 17 |
| Figure S1. PCoA analysis of 6,889 loci. | 18 |
| Figure S2. Schematic of “sec_contact_asym_mig_three_epoch” model. | 19 |
| Figure S3. Parsimony network of mtDNA haplotypes. | 20 |
| Figure S4. δaδi results. | 21 |
| Figure S5. Google Earth map of 12 km density trough. | 22 |
| Supplemental materials and methods | 23– 26 |

**Table S1**. Details of all individuals used in the study, including if the individual was sequenced (SNPs and/or mitochondrial ND4 gene), had dorsal/ventral photos and spectral reflectance measurements, and the sex (M = male, F = female). Population [Southern: Kangaroo Island (KI), Mainland South (MS); Northern: Northern Flinders Ranges (NFR), Southern Flinders Ranges (SFR), Olary Ranges (OR); and Contact Zone (CZ)], year, and cline site (if applicable) are shown.

| ID | SNPs | ND4 | Photos | Specs | Sex | Pop | Year | Cline Site | Latitude | Longitude |
|---|---|---|---|---|---|---|---|---|---|---|
| KI206 | Y | N | N | N | M | KI | 2010 | NA | -35.965 | 136.653 |
| KI208 | Y | N | N | N | M | KI | 2011 | NA | -35.965 | 136.653 |
| KI194 | Y | N | N | N | F | KI | 2010 | NA | -35.964 | 136.653 |
| KI179 | Y | N | N | N | M | KI | 2010 | NA | -35.963 | 136.654 |
| KI193 | Y | N | N | N | M | KI | 2010 | NA | -35.963 | 136.654 |
| KI181 | Y | N | N | N | M | KI | 2010 | NA | -35.960 | 136.655 |
| KI215 | Y | N | N | N | M | KI | 2011 | NA | -35.621 | 137.210 |
| R45184 | Y | N | N | N | NA | MS | 1994 | NA | -35.642 | 138.329 |
| R49234 | Y | N | N | N | NA | MS | 1996 | NA | -35.601 | 138.279 |
| R53785 | Y | N | N | N | F | MS | 2000 | NA | -35.194 | 138.919 |
| R55041 | Y | N | N | N | F | MS | 2000 | NA | -35.160 | 138.578 |
| M232 | Y | N | N | N | M | MS | 2011 | NA | -34.908 | 138.707 |
| M235 | Y | N | N | N | M | MS | 2011 | NA | -34.906 | 138.702 |
| M248 | Y | N | N | N | M | MS | 2011 | NA | -34.906 | 138.707 |
| R42978 | Y | N | N | N | NA | MS | 1993 | NA | -34.683 | 138.850 |
| KS630 | Y | N | N | N | M | MS | 2012 | NA | -34.582 | 139.015 |
| KS631 | Y | N | N | N | M | MS | 2012 | NA | -34.582 | 139.015 |
| KS632 | Y | N | N | N | M | MS | 2012 | NA | -34.582 | 139.015 |
| KS634 | Y | N | N | N | F | MS | 2012 | NA | -34.581 | 139.015 |
| R27416 | N | Y | N | N | NA | MS | 2012 | 1 | -34.842 | 139.133 |
| 638 | N | Y | N | N | F | MS | 2012 | 1 | -34.831 | 139.130 |
| 639 | N | Y | N | N | F | MS | 2012 | 1 | -34.494 | 139.201 |
| 640 | N | Y | N | N | F | MS | 2012 | 1 | -34.830 | 139.129 |
| 641 | N | Y | N | N | M | MS | 2012 | 1 | -34.494 | 139.201 |
| 642 | N | Y | N | N | F | MS | 2012 | 1 | -34.833 | 139.128 |
| PB1 | Y | N | Y | N | M | MS | 2013 | 1 | -34.834 | 139.128 |
| S42 | Y | N | Y | Y | M | MS | 2016 | 1 | -34.830 | 139.129 |
| S32 | N | N | Y | Y | M | MS | 2015 | 1 | -34.830 | 139.128 |
| S44 | N | N | Y | Y | M | MS | 2016 | 1 | -34.829 | 139.129 |
| S31 | Y | Y | Y | Y | M | MS | 2015 | 1 | -34.829 | 139.130 |
| S39 | N | N | Y | Y | M | MS | 2015 | 1 | -34.829 | 139.129 |
| S47 | Y | Y | Y | Y | M | MS | 2016 | 1 | -34.828 | 139.124 |
| S30 | N | N | Y | Y | M | MS | 2015 | 1 | -34.826 | 139.129 |
| S40 | N | N | Y | Y | M | MS | 2015 | 1 | -34.826 | 139.160 |
| S33 | N | N | Y | Y | M | MS | 2015 | 1 | -34.822 | 139.162 |
| S29 | Y | N | Y | Y | M | MS | 2015 | 1 | -34.822 | 139.116 |
| S8 | Y | Y | N | N | F | MS | 2015 | 1 | -34.822 | 139.116 |

| | | | | | | | | | | |
|---|---|---|---|---|---|---|---|---|---|---|
| S27 | N | N | Y | Y | M | MS | 2015 | 1 | -34.821 | 139.121 |
| S28 | N | N | Y | Y | M | MS | 2015 | 1 | -34.821 | 139.121 |
| PB9 | N | N | Y | N | M | MS | 2013 | 1 | -34.821 | 139.160 |
| PB10 | N | N | Y | N | M | MS | 2013 | 1 | -34.821 | 139.160 |
| PB4 | N | N | Y | N | M | MS | 2013 | 1 | -34.820 | 139.137 |
| PB8 | Y | Y | Y | N | M | MS | 2013 | 1 | -34.821 | 139.160 |
| 641 | N | N | Y | Y | M | MS | 2012 | 1 | -34.494 | 139.201 |
| PB2 | N | N | Y | N | M | MS | 2013 | 1 | -34.820 | 139.160 |
| PB3 | N | N | Y | N | M | MS | 2013 | 1 | -34.820 | 139.132 |
| PB5 | N | N | Y | N | M | MS | 2013 | 1 | -34.819 | 139.162 |
| S21 | Y | N | Y | Y | M | MS | 2015 | 1 | -34.818 | 139.162 |
| PB6 | N | N | Y | N | M | MS | 2013 | 1 | -34.818 | 139.163 |
| S36 | N | N | Y | Y | M | MS | 2015 | 1 | -34.818 | 139.163 |
| PB7 | N | N | Y | N | M | MS | 2013 | 1 | -34.818 | 139.164 |
| S26 | N | N | Y | Y | M | MS | 2015 | 1 | -34.817 | 139.117 |
| 710 | Y | N | N | N | M | MS | 2013 | 1 | -34.817 | 139.117 |
| S25 | Y | N | Y | Y | M | MS | 2015 | 1 | -34.817 | 139.116 |
| S24 | N | N | Y | Y | M | MS | 2015 | 1 | -34.817 | 139.164 |
| S35 | N | N | Y | Y | M | MS | 2015 | 1 | -34.817 | 139.114 |
| S22 | N | N | Y | Y | M | MS | 2015 | 1 | -34.817 | 139.165 |
| S23 | N | N | Y | Y | M | MS | 2015 | 1 | -34.817 | 139.165 |
| S37 | N | N | Y | Y | M | MS | 2015 | 1 | -34.816 | 139.163 |
| S38 | N | N | Y | Y | M | MS | 2015 | 1 | -33.022 | 138.000 |
| 1142 | N | N | Y | Y | M | MS | 2017 | 2 | -34.793 | 139.203 |
| 1179 | N | N | Y | Y | M | MS | 2017 | 2 | -34.742 | 139.212 |
| 1177 | Y | N | Y | Y | M | MS | 2017 | 2 | -34.742 | 139.211 |
| 1178 | N | N | Y | Y | M | MS | 2017 | 2 | -34.740 | 139.211 |
| 1145 | Y | N | Y | Y | M | MS | 2017 | 2 | -34.726 | 139.214 |
| 1146 | Y | N | Y | Y | M | MS | 2017 | 2 | -34.725 | 139.214 |
| 703 | Y | N | Y | N | M | MS | 2013 | 2 | -34.725 | 139.214 |
| 1140 | N | N | Y | Y | M | MS | 2017 | 2 | -34.723 | 139.216 |
| 1143 | N | N | Y | Y | M | MS | 2017 | 2 | -34.723 | 139.214 |
| 1141 | Y | N | Y | Y | M | MS | 2017 | 2 | -34.723 | 139.215 |
| 1144 | Y | N | Y | Y | M | MS | 2017 | 2 | -34.723 | 139.215 |
| 1171 | N | N | Y | Y | M | MS | 2017 | 2 | -34.721 | 139.215 |
| 1164 | N | N | Y | Y | M | MS | 2017 | 2 | -34.721 | 139.215 |
| 1167 | N | N | Y | Y | M | MS | 2017 | 2 | -34.721 | 139.213 |
| 1168 | Y | N | Y | Y | M | MS | 2017 | 2 | -34.721 | 139.213 |
| 1169 | N | N | Y | Y | M | MS | 2017 | 2 | -34.721 | 139.213 |
| 1170 | N | N | Y | Y | M | MS | 2017 | 2 | -34.721 | 139.213 |
| 1166 | N | N | Y | Y | M | MS | 2017 | 2 | -34.720 | 139.213 |
| 709 | Y | Y | N | N | M | MS | 2013 | 2 | -34.720 | 139.207 |
| 700 | N | N | Y | N | M | MS | 2013 | 2 | -34.720 | 139.214 |
| 1165 | Y | N | Y | Y | M | MS | 2017 | 2 | -34.720 | 139.214 |
| 706 | Y | N | N | N | M | MS | 2013 | 2 | -34.718 | 139.206 |
| S1107 | Y | N | Y | Y | M | MS | 2016 | 2 | -34.701 | 139.206 |
| 1180 | N | N | Y | Y | M | CZ | 2017 | 3 | -34.591 | 139.218 |
| H1106 | Y | N | Y | Y | M | CZ | 2016 | 3 | -34.590 | 139.218 |

| | | | | | | | | | | |
|---|---|---|---|---|---|---|---|---|---|---|
| 1186 | N | N | Y | Y | M | CZ | 2017 | 3 | -34.590 | 139.209 |
| 1185 | Y | Y | Y | Y | M | CZ | 2017 | 3 | -34.589 | 139.211 |
| 1187 | N | N | Y | Y | M | CZ | 2017 | 3 | -34.588 | 139.215 |
| 1184 | N | N | Y | Y | M | CZ | 2017 | 3 | -34.588 | 139.215 |
| H1104 | Y | Y | N | N | F | CZ | 2016 | 3 | -34.588 | 139.218 |
| 1182 | N | N | Y | Y | M | CZ | 2017 | 3 | -34.588 | 139.217 |
| 1183 | N | N | Y | Y | M | CZ | 2017 | 3 | -34.587 | 139.216 |
| 759 | Y | N | Y | N | M | CZ | 2013 | 3 | -34.587 | 139.207 |
| 758 | N | N | Y | N | M | CZ | 2013 | 3 | -34.587 | 139.206 |
| 756 | Y | Y | Y | N | M | CZ | 2013 | 3 | -34.586 | 139.206 |
| 754 | Y | N | N | N | M | CZ | 2013 | 3 | -34.586 | 139.205 |
| 1215 | N | N | Y | Y | M | CZ | 2017 | 3 | -34.585 | 139.216 |
| 1217 | N | N | Y | Y | M | CZ | 2017 | 3 | -34.585 | 139.216 |
| 1216 | N | N | Y | Y | M | CZ | 2017 | 3 | -34.585 | 139.216 |
| H1100 | Y | N | Y | Y | M | CZ | 2016 | 3 | -34.583 | 139.215 |
| H1101 | Y | N | Y | Y | M | CZ | 2016 | 3 | -34.583 | 139.215 |
| H1102 | Y | Y | N | N | F | CZ | 2016 | 3 | -34.579 | 139.215 |
| H1103 | Y | Y | N | N | F | CZ | 2016 | 3 | -34.577 | 139.215 |
| H1099 | Y | N | Y | Y | M | CZ | 2016 | 3 | -34.577 | 139.203 |
| H1021 | Y | N | Y | Y | M | CZ | 2015 | 4 | -34.571 | 139.205 |
| 739 | Y | N | Y | N | M | CZ | 2013 | 4 | -34.564 | 139.211 |
| 738 | Y | Y | N | N | F | CZ | 2013 | 4 | -34.563 | 139.208 |
| 740 | Y | N | Y | N | M | CZ | 2013 | 4 | -34.563 | 139.213 |
| 741 | Y | Y | N | N | M | CZ | 2013 | 4 | -34.563 | 139.212 |
| 737 | Y | N | Y | N | M | CZ | 2013 | 4 | -34.563 | 139.207 |
| 753 | N | N | Y | N | M | CZ | 2013 | 4 | -34.561 | 139.193 |
| 716 | Y | N | Y | N | M | CZ | 2013 | 4 | -34.561 | 139.202 |
| 712 | N | N | Y | N | M | CZ | 2013 | 4 | -34.561 | 139.203 |
| 713 | Y | N | N | N | M | CZ | 2013 | 4 | -34.559 | 139.205 |
| H1020 | Y | N | Y | Y | M | CZ | 2015 | 4 | -34.558 | 139.202 |
| H1019 | N | N | Y | Y | M | CZ | 2015 | 4 | -34.558 | 139.203 |
| H1009 | N | N | Y | Y | M | CZ | 2015 | 4 | -34.558 | 139.177 |
| H1018 | N | N | Y | Y | M | CZ | 2015 | 4 | -34.557 | 139.204 |
| 715 | N | N | Y | N | M | CZ | 2013 | 4 | -34.557 | 139.204 |
| H1017 | Y | Y | Y | Y | M | CZ | 2015 | 4 | -34.557 | 139.205 |
| H1051 | Y | N | Y | Y | M | CZ | 2016 | 4 | -34.557 | 139.200 |
| H1039 | N | N | Y | Y | M | CZ | 2015 | 4 | -34.556 | 139.199 |
| H1007 | Y | N | Y | Y | M | CZ | 2015 | 4 | -34.556 | 139.200 |
| H1048 | N | N | Y | Y | M | CZ | 2015 | 4 | -34.556 | 139.199 |
| H1008 | Y | N | Y | Y | M | CZ | 2015 | 4 | -34.556 | 139.198 |
| H1046 | N | N | Y | Y | M | CZ | 2015 | 4 | -34.556 | 139.198 |
| 746 | N | N | Y | N | M | CZ | 2013 | 4 | -34.556 | 139.199 |
| 742 | Y | Y | N | N | F | CZ | 2013 | 4 | -34.556 | 139.198 |
| H1049 | Y | N | Y | Y | M | CZ | 2016 | 4 | -34.556 | 139.199 |
| 743 | N | N | Y | N | M | CZ | 2013 | 4 | -34.556 | 139.198 |
| 745 | N | N | Y | N | M | CZ | 2013 | 4 | -34.556 | 139.197 |
| H1045 | N | N | Y | Y | M | CZ | 2015 | 4 | -34.556 | 139.199 |
| H1052 | Y | N | N | N | F | CZ | 2016 | 4 | -34.556 | 139.200 |

| | | | | | | | | | | |
|---|---|---|---|---|---|---|---|---|---|---|
| H1050 | Y | N | Y | Y | M | CZ | 2016 | 4 | -34.556 | 139.200 |
| H1044 | N | N | Y | Y | M | CZ | 2015 | 4 | -34.556 | 139.200 |
| H1055 | Y | Y | N | N | F | CZ | 2016 | 4 | -34.556 | 139.201 |
| 744 | N | N | Y | N | M | CZ | 2013 | 4 | -34.556 | 139.202 |
| H1053 | Y | Y | Y | Y | M | CZ | 2016 | 4 | -34.556 | 139.200 |
| H1054 | Y | N | Y | Y | M | CZ | 2016 | 4 | -34.556 | 139.202 |
| H1056 | Y | Y | N | N | F | CZ | 2016 | 4 | -34.556 | 139.202 |
| H1010 | N | N | Y | Y | M | CZ | 2015 | 4 | -34.556 | 139.203 |
| H1014 | Y | N | Y | Y | M | CZ | 2015 | 4 | -34.556 | 139.203 |
| H1016 | Y | Y | Y | Y | M | CZ | 2015 | 4 | -34.556 | 139.202 |
| H1022 | Y | Y | Y | Y | M | CZ | 2015 | 4 | -34.556 | 139.201 |
| 748 | N | N | Y | N | M | CZ | 2013 | 4 | -34.556 | 139.199 |
| H1011 | Y | N | Y | Y | M | CZ | 2015 | 4 | -34.555 | 139.203 |
| H1015 | N | N | Y | Y | M | CZ | 2015 | 4 | -34.555 | 137.203 |
| H1013 | Y | N | Y | Y | M | CZ | 2015 | 4 | -34.555 | 139.203 |
| H1012 | Y | N | Y | Y | M | CZ | 2015 | 4 | -34.555 | 139.203 |
| H1023 | N | N | Y | Y | M | CZ | 2015 | 4 | -34.555 | 139.202 |
| H1027 | Y | Y | Y | Y | M | CZ | 2015 | 4 | -34.555 | 139.206 |
| H1041 | N | N | Y | Y | M | CZ | 2015 | 4 | -34.555 | 139.205 |
| 750 | N | N | Y | N | M | CZ | 2013 | 4 | -34.555 | 139.202 |
| H1042 | N | N | Y | Y | M | CZ | 2015 | 4 | -34.555 | 139.204 |
| H1043 | N | N | Y | Y | M | CZ | 2015 | 4 | -34.555 | 139.204 |
| H1040 | N | N | Y | Y | M | CZ | 2015 | 4 | -34.555 | 139.204 |
| H1029 | N | N | Y | Y | M | CZ | 2015 | 4 | -34.555 | 139.204 |
| H1025 | Y | N | Y | Y | M | CZ | 2015 | 4 | -34.555 | 139.207 |
| H1026 | N | N | Y | Y | M | CZ | 2015 | 4 | -34.554 | 139.207 |
| H1024 | N | N | Y | Y | M | CZ | 2015 | 4 | -34.554 | 139.202 |
| H1028 | N | N | Y | Y | M | CZ | 2015 | 4 | -34.554 | 139.206 |
| 751 | N | N | Y | N | M | CZ | 2013 | 4 | -34.554 | 139.202 |
| 749 | Y | Y | N | N | F | CZ | 2013 | 4 | -34.552 | 139.203 |
| H1086 | Y | N | Y | Y | M | CZ | 2016 | 4 | -34.551 | 139.192 |
| H1089 | Y | N | Y | Y | M | CZ | 2016 | 5 | -34.542 | 139.197 |
| H1091 | Y | Y | Y | Y | M | CZ | 2016 | 5 | -34.540 | 139.202 |
| H1092 | Y | Y | Y | Y | M | CZ | 2016 | 5 | -34.540 | 139.202 |
| H1057 | Y | N | Y | Y | M | CZ | 2016 | 5 | -34.540 | 139.198 |
| H1059 | Y | Y | Y | Y | M | CZ | 2016 | 5 | -34.539 | 139.196 |
| H1090 | Y | N | Y | Y | M | CZ | 2016 | 5 | -34.539 | 139.194 |
| 636 | Y | N | N | N | M | CZ | 2012 | 5 | -34.537 | 138.999 |
| H1097 | Y | Y | Y | Y | M | CZ | 2016 | 5 | -34.534 | 139.198 |
| H1060 | Y | Y | Y | Y | M | CZ | 2016 | 5 | -34.531 | 139.206 |
| H1093 | N | N | Y | Y | M | CZ | 2016 | 5 | -34.528 | 139.197 |
| H1096 | Y | N | Y | Y | M | CZ | 2016 | 5 | -34.526 | 139.195 |
| H1098 | Y | Y | Y | Y | M | CZ | 2016 | 5 | -34.526 | 139.195 |
| 637 | N | Y | N | N | F | CZ | 2012 | 6 | -34.494 | 139.201 |
| H1084 | Y | N | Y | Y | M | CZ | 2016 | 6 | -34.515 | 139.200 |
| H1075 | Y | N | N | N | F | CZ | 2016 | 6 | -34.513 | 139.201 |
| H1076 | N | N | Y | Y | M | CZ | 2016 | 6 | -34.513 | 139.201 |
| H1082 | Y | Y | Y | Y | M | CZ | 2016 | 6 | -34.513 | 139.200 |

| | | | | | | | | | | |
|---|---|---|---|---|---|---|---|---|---|---|
| H1078 | Y | N | Y | Y | M | CZ | 2016 | 6 | -34.513 | 139.200 |
| H1079 | Y | N | Y | Y | M | CZ | 2016 | 6 | -34.508 | 139.202 |
| H1081 | Y | N | Y | Y | M | CZ | 2016 | 6 | -34.507 | 139.202 |
| H1072 | N | N | Y | Y | M | CZ | 2016 | 6 | -34.503 | 139.205 |
| H1074 | Y | N | Y | Y | M | CZ | 2016 | 6 | -34.503 | 139.205 |
| H1071 | Y | Y | Y | Y | M | CZ | 2016 | 6 | -34.498 | 139.201 |
| H1068 | Y | N | Y | Y | M | CZ | 2016 | 6 | -34.495 | 139.202 |
| H1069 | N | N | Y | Y | M | CZ | 2016 | 6 | -34.495 | 139.202 |
| H1064 | Y | N | Y | Y | M | CZ | 2016 | 6 | -34.494 | 139.201 |
| H1065 | N | N | Y | Y | M | CZ | 2016 | 6 | -34.494 | 139.202 |
| 1218 | N | N | Y | Y | M | CZ | 2017 | 6 | -34.493 | 139.202 |
| 1220 | Y | N | Y | Y | M | CZ | 2017 | 6 | -34.492 | 139.203 |
| H1067 | Y | N | Y | Y | M | CZ | 2016 | 6 | -34.490 | 139.203 |
| 1222 | N | N | Y | Y | M | CZ | 2017 | 6 | -34.494 | 139.200 |
| 1221 | N | N | Y | Y | M | CZ | 2017 | 6 | -34.494 | 139.201 |
| 1223 | Y | N | Y | Y | M | CZ | 2017 | 7 | 34.494 | 139.201 |
| H1108 | Y | N | N | N | M | CZ | 2016 | 7 | -34.465 | 139.203 |
| H1109 | Y | N | Y | Y | M | CZ | 2016 | 7 | -34.464 | 139.203 |
| H1111 | Y | Y | Y | Y | M | CZ | 2016 | 7 | -34.451 | 139.218 |
| H1110 | Y | N | N | N | F | CZ | 2016 | 7 | -34.451 | 139.219 |
| H1113 | Y | N | Y | Y | M | CZ | 2016 | 7 | -34.449 | 139.215 |
| H1112 | Y | N | Y | Y | M | CZ | 2016 | 7 | -34.448 | 139.209 |
| H1114 | Y | N | Y | Y | M | CZ | 2016 | 7 | -34.443 | 39.210 |
| H1115 | Y | N | N | N | F | CZ | 2016 | 7 | -34.436 | 139.212 |
| 627 | N | Y | N | N | M | CZ | 2012 | 8 | -34.411 | 139.159 |
| 628 | N | Y | N | N | F | CZ | 2012 | 8 | -34.411 | 139.159 |
| 629 | N | Y | N | N | F | CZ | 2012 | 8 | -34.413 | 139.161 |
| 719 | Y | Y | N | N | M | CZ | 2013 | 8 | -34.421 | 139.187 |
| 721 | Y | N | N | N | M | CZ | 2013 | 8 | -34.414 | 139.168 |
| 728 | Y | N | N | N | M | CZ | 2013 | 8 | -34.412 | 139.164 |
| 720 | Y | N | N | N | M | CZ | 2013 | 8 | -34.411 | 139.158 |
| H1139 | Y | N | Y | Y | M | CZ | 2016 | 8 | -34.409 | 139.210 |
| H1136 | Y | N | N | N | F | CZ | 2016 | 8 | -34.408 | 139.215 |
| H1137 | Y | N | Y | Y | M | CZ | 2016 | 8 | -34.408 | 139.215 |
| H1117 | Y | N | Y | Y | M | OR | 2016 | 9 | -34.366 | 139.164 |
| H1116 | Y | Y | Y | Y | M | OR | 2016 | 9 | -34.364 | 139.160 |
| 1173 | Y | N | Y | Y | M | OR | 2017 | 9 | -34.360 | 139.208 |
| 1174 | N | N | Y | Y | M | OR | 2017 | 9 | -34.360 | 139.208 |
| 1175 | Y | N | Y | Y | M | OR | 2017 | 9 | -34.358 | 139.209 |
| 1210 | N | N | Y | Y | M | OR | 2017 | 9 | -34.351 | 139.190 |
| 1224 | Y | N | Y | Y | M | OR | 2017 | 9 | -34.350 | 139.188 |
| 1225 | N | N | Y | Y | M | OR | 2017 | 9 | -34.350 | 139.188 |
| 1211 | N | N | Y | Y | M | OR | 2017 | 9 | -34.350 | 139.188 |
| 1209 | N | N | Y | Y | M | OR | 2017 | 9 | -34.349 | 139.189 |
| 1208 | Y | N | Y | Y | M | OR | 2017 | 9 | -34.349 | 139.188 |
| 1212 | Y | Y | Y | Y | M | OR | 2017 | 9 | -34.348 | 139.189 |
| 1214 | Y | N | Y | Y | M | OR | 2017 | 9 | -34.348 | 139.189 |
| 1213 | Y | N | Y | Y | M | OR | 2017 | 9 | -34.348 | 139.189 |

| | | | | | | | | | | |
|---|---|---|---|---|---|---|---|---|---|---|
| 1159 | N | N | Y | Y | M | OR | 2017 | 9 | -34.332 | 139.173 |
| 1176 | N | N | Y | Y | M | OR | 2017 | 9 | -34.332 | 139.172 |
| 1163 | Y | N | Y | Y | M | OR | 2017 | 9 | -34.332 | 139.170 |
| 1162 | Y | Y | Y | Y | M | OR | 2017 | 9 | -34.331 | 139.173 |
| 1160 | Y | N | Y | Y | M | OR | 2017 | 9 | -34.331 | 139.174 |
| 1161 | Y | Y | N | N | F | OR | 2017 | 9 | -34.331 | 139.174 |
| H1121 | N | N | Y | Y | M | OR | 2016 | 9 | -34.331 | 139.174 |
| 1158 | Y | N | Y | Y | M | OR | 2017 | 9 | -34.331 | 139.174 |
| H1122 | Y | N | Y | Y | M | OR | 2016 | 9 | -34.331 | 139.173 |
| H1123 | Y | Y | Y | Y | M | OR | 2016 | 9 | -34.331 | 139.174 |
| H1124 | Y | N | Y | Y | M | OR | 2016 | 10 | -34.231 | 139.118 |
| H1125 | Y | Y | Y | Y | M | OR | 2016 | 10 | -34.228 | 139.118 |
| 1207 | Y | N | Y | Y | M | OR | 2017 | 11 | -34.167 | 139.069 |
| H1133 | Y | N | Y | Y | M | OR | 2016 | 11 | -34.167 | 139.069 |
| 1206 | N | N | Y | Y | M | OR | 2017 | 11 | -34.167 | 139.069 |
| 1156 | Y | N | Y | Y | M | OR | 2017 | 11 | -34.166 | 139.070 |
| 1155 | Y | N | Y | Y | M | OR | 2017 | 11 | -34.166 | 139.071 |
| 1152 | Y | Y | Y | Y | M | OR | 2017 | 11 | -34.132 | 139.048 |
| 1148 | Y | N | Y | Y | M | OR | 2017 | 11 | -34.128 | 139.048 |
| 1154 | Y | N | Y | Y | M | OR | 2017 | 11 | -34.127 | 139.048 |
| 1149 | Y | Y | N | N | F | OR | 2017 | 11 | -34.127 | 139.048 |
| 1151 | N | N | Y | Y | M | OR | 2017 | 11 | -34.127 | 139.048 |
| H1131 | Y | Y | Y | Y | M | OR | 2016 | 11 | -34.127 | 139.049 |
| 1153 | N | N | Y | Y | M | OR | 2017 | 11 | -34.125 | 139.047 |
| H1135 | Y | N | Y | Y | M | OR | 2016 | 12 | -34.091 | 139.035 |
| 626 | Y | Y | N | N | M | OR | 2012 | 12 | -34.070 | 138.957 |
| R58318 | Y | Y | N | N | NA | OR | 2003 | 12 | -34.031 | 138.935 |
| N27 | Y | N | Y | Y | M | OR | 2015 | 12 | -33.935 | 138.952 |
| N16 | Y | N | Y | Y | M | OR | 2015 | 12 | -33.934 | 138.952 |
| 623 | Y | N | N | N | M | OR | 2012 | 12 | -33.934 | 138.952 |
| 618 | Y | Y | N | N | M | OR | 2012 | 12 | -33.934 | 138.952 |
| R26108 | Y | N | N | N | NA | OR | 1984 | 12 | -33.917 | 138.550 |
| 619 | N | Y | N | N | M | OR | 2012 | 12 | -33.934 | 138.952 |
| 620 | N | Y | N | N | F | OR | 2012 | 12 | -33.934 | 138.952 |
| 622 | N | Y | N | N | F | OR | 2012 | 12 | -33.934 | 138.952 |
| 624 | N | Y | N | N | F | OR | 2012 | 12 | - | - |
| 625 | N | Y | N | N | F | OR | 2012 | 12 | -34.070 | 138.957 |
| N28 | N | N | Y | Y | M | OR | 2015 | 12 | -33.934 | 138.953 |
| 623 | Y | N | N | N | NA | OR | 1992 | 12 | -33.934 | 138.952 |
| R40708 | Y | Y | N | N | NA | OR | 1992 | 13 | -33.667 | 138.950 |
| 1205 | N | N | Y | Y | M | OR | 2017 | 13 | -33.662 | 139.042 |
| 1197 | N | N | Y | Y | M | OR | 2017 | 13 | -33.631 | 139.004 |
| 1195 | Y | Y | Y | Y | M | OR | 2017 | 13 | -33.631 | 139.004 |
| 1203 | N | N | Y | Y | M | OR | 2017 | 13 | -33.628 | 138.996 |
| 1194 | N | N | Y | Y | M | OR | 2017 | 13 | -33.628 | 139.003 |
| 1190 | N | Y | Y | Y | M | OR | 2017 | 13 | -33.628 | 139.001 |
| 1189 | Y | Y | Y | Y | M | OR | 2017 | 13 | -33.628 | 138.999 |
| 1193 | N | N | Y | Y | M | OR | 2017 | 13 | -33.628 | 139.002 |

| | | | | | | | | | | |
|---|---|---|---|---|---|---|---|---|---|---|
| 1188 | N | N | Y | Y | M | OR | 2017 | 13 | -33.627 | 138.999 |
| 1191 | Y | N | Y | Y | M | OR | 2017 | 13 | -33.627 | 138.994 |
| 1200 | Y | Y | Y | Y | M | OR | 2017 | 13 | -33.615 | 138.996 |
| 1201 | N | N | Y | Y | M | OR | 2017 | 13 | -33.615 | 138.996 |
| 1202 | N | N | Y | Y | M | OR | 2017 | 13 | -33.615 | 138.996 |
| R41631 | N | Y | N | N | M | OR | 1992 | 13 | -33.675 | 138.961 |
| R38012 | N | Y | N | N | NA | OR | 1991 | 13 | -33.833 | 139.017 |
| 91 | N | Y | N | N | M | OR | 2010 | 13 | -33.835 | 139.034 |
| N32 | N | N | Y | Y | M | OR | 2015 | 14 | -33.444 | 139.102 |
| N36 | Y | N | Y | Y | M | OR | 2015 | 14 | -33.442 | 139.104 |
| N34 | N | N | Y | Y | M | OR | 2015 | 14 | -33.442 | 139.104 |
| N30 | Y | Y | Y | Y | M | OR | 2015 | 14 | -33.441 | 139.103 |
| N33 | N | N | Y | Y | M | OR | 2015 | 14 | -33.441 | 139.103 |
| N42 | N | N | Y | Y | M | OR | 2016 | 14 | -33.439 | 139.103 |
| N49 | N | N | Y | Y | M | OR | 2016 | 14 | -33.439 | 139.101 |
| N44 | Y | Y | N | N | F | OR | 2016 | 14 | -33.439 | 139.101 |
| N21 | Y | N | Y | Y | M | OR | 2015 | 14 | -33.411 | 139.111 |
| N40 | Y | Y | Y | Y | M | OR | 2015 | 14 | -33.411 | 139.111 |
| N25 | Y | Y | Y | Y | M | OR | 2015 | 14 | -33.410 | 139.095 |
| N24 | N | N | Y | Y | M | OR | 2015 | 14 | -33.410 | 139.096 |
| N29 | N | N | Y | Y | M | OR | 2015 | 14 | -33.411 | 139.112 |
| N39 | N | N | Y | Y | M | OR | 2015 | 14 | -33.411 | 139.111 |
| N26 | N | N | Y | Y | M | OR | 2015 | 14 | -33.411 | 139.102 |
| N22 | N | N | Y | Y | M | OR | 2015 | 14 | -33.411 | 139.103 |
| N23 | N | N | Y | Y | M | OR | 2015 | 14 | -33.412 | 139.098 |
| N37 | N | N | Y | Y | M | OR | 2015 | 14 | -33.442 | 139.101 |
| N31 | N | N | Y | Y | M | OR | 2015 | 14 | -33.442 | 139.107 |
| N38 | N | N | Y | Y | M | OR | 2015 | 14 | -33.444 | 139.103 |
| R58759 | N | Y | N | N | M | OR | 2004 | 14 | -33.434 | 139.079 |
| R53529 | N | Y | N | N | NA | OR | 1999 | 14 | -33.345 | 139.261 |
| R41362 | Y | N | N | N | NA | OR | 1992 | NA | -33.350 | 139.103 |
| R56417 | Y | N | N | N | NA | OR | 2001 | NA | -33.234 | 138.250 |
| R41167 | Y | N | N | N | M | OR | 1992 | NA | -33.200 | 139.619 |
| R41638 | Y | N | N | N | F | OR | 1992 | NA | -33.124 | 139.196 |
| R41771 | Y | N | N | N | NA | OR | 1992 | NA | -33.061 | 139.831 |
| BS377 | Y | N | N | N | F | OR | 2011 | NA | -32.095 | 140.274 |
| BS378 | Y | N | N | N | M | OR | 2011 | NA | -32.095 | 140.274 |
| BS379 | Y | N | N | N | M | OR | 2011 | NA | -32.094 | 140.274 |
| R60607 | Y | N | N | N | F | OR | 2005 | NA | -32.067 | 140.333 |
| BS398 | Y | N | N | N | M | OR | 2011 | NA | -31.981 | 140.208 |
| BS400 | Y | N | N | N | F | OR | 2011 | NA | -31.981 | 140.210 |
| TG02 | Y | N | N | N | M | SFR | 2015 | NA | - | - |
| TGG1 | Y | N | N | N | M | SFR | 2015 | NA | - | - |
| TGO1 | Y | N | N | N | M | SFR | 2015 | NA | - | - |
| TGOY1 | Y | N | N | N | M | SFR | 2015 | NA | - | - |
| TGY1 | Y | N | N | N | M | SFR | 2015 | NA | - | - |
| TG_F8 | Y | N | N | N | F | SFR | 2010 | NA | - | - |
| R31224 | Y | N | N | N | NA | SFR | 1987 | NA | -32.867 | 139.933 |

| | | | | | | | | | | |
|---|---|---|---|---|---|---|---|---|---|---|
| MR595 | Y | N | N | N | M | SFR | 2012 | NA | -32.843 | 138.053 |
| MR588 | Y | N | N | N | M | SFR | 2012 | NA | -32.842 | 138.057 |
| MR590 | Y | N | N | N | M | SFR | 2012 | NA | -32.842 | 138.057 |
| MR591 | Y | N | N | N | M | SFR | 2012 | NA | -32.842 | 138.057 |
| AG63 | Y | N | N | N | F | SFR | 2010 | NA | -32.745 | 138.073 |
| R41137 | Y | N | N | N | NA | SFR | 1992 | NA | -32.592 | 139.861 |
| DP88 | Y | N | N | N | M | SFR | 2010 | NA | -32.416 | 137.993 |
| DS141 | Y | N | N | N | F | SFR | 2010 | NA | -32.310 | 137.980 |
| WG136 | Y | N | N | N | F | SFR | 2010 | NA | -32.188 | 138.011 |
| WG143 | Y | N | N | N | M | SFR | 2010 | NA | -32.188 | 138.010 |
| WG166 | Y | N | N | N | M | SFR | 2010 | NA | -32.188 | 138.009 |
| WG80 | Y | N | N | N | M | SFR | 2010 | NA | -32.188 | 138.009 |
| WG135 | Y | N | N | N | F | SFR | 2010 | NA | -32.188 | 138.011 |
| WG146 | Y | N | N | N | M | SFR | 2010 | NA | -32.187 | 138.008 |
| WG145 | Y | N | N | N | M | SFR | 2010 | NA | -32.186 | 138.008 |
| YC58 | Y | N | N | N | NA | SFR | 2010 | NA | -31.958 | 138.377 |
| YC59 | Y | N | N | N | M | SFR | 2010 | NA | -31.958 | 138.377 |
| YC60 | Y | N | N | N | M | SFR | 2010 | NA | -31.956 | 138.372 |
| YC127 | Y | N | N | N | M | SFR | 2010 | NA | -31.956 | 138.372 |
| YC93 | Y | N | N | N | M | SFR | 2010 | NA | -31.956 | 138.372 |
| YC110 | Y | N | N | N | M | SFR | 2010 | NA | -31.954 | 138.375 |
| YC104 | Y | N | N | N | M | SFR | 2010 | NA | -31.953 | 138.374 |
| YC105 | Y | N | N | N | F | SFR | 2010 | NA | -31.953 | 138.374 |
| R57145 | Y | N | N | N | M | SFR | - | NA | -31.938 | 138.105 |
| R60917 | Y | N | N | N | NA | SFR | 2005 | NA | -31.625 | 138.594 |
| W_M31 | Y | N | N | N | M | SFR | 2010 | NA | -31.546 | 138.603 |
| W_M23 | Y | N | N | N | M | SFR | 2010 | NA | -31.538 | 138.599 |
| W_M22 | Y | N | N | N | M | SFR | 2010 | NA | -31.538 | 138.599 |
| BG51 | Y | N | N | N | M | SFR | 2010 | NA | -31.418 | 138.558 |
| R52282 | Y | N | N | N | NA | SFR | 1999 | NA | -31.395 | 138.811 |
| A535 | Y | N | N | N | M | SFR | 2012 | NA | -31.281 | 138.582 |
| A529 | Y | N | N | N | M | SFR | 2012 | NA | -31.280 | 138.583 |
| A504 | Y | N | N | N | M | SFR | 2012 | NA | -31.280 | 138.584 |
| A510 | Y | N | N | N | F | SFR | 2012 | NA | -31.279 | 138.591 |
| A518 | Y | N | N | N | M | SFR | 2012 | NA | -31.279 | 138.593 |
| A47 | Y | N | N | N | M | SFR | 2010 | NA | -31.277 | 138.599 |
| R52684 | Y | N | N | N | NA | NFR | 1999 | NA | -30.768 | 138.707 |
| R51819 | Y | N | N | N | M | NFR | 1998 | NA | -30.611 | 138.803 |
| R52317 | Y | N | N | N | NA | NFR | 1999 | NA | -30.598 | 139.133 |
| R52909 | Y | N | N | N | NA | NFR | 1999 | NA | -30.428 | 139.145 |
| R31653 | Y | N | N | N | NA | NFR | 1987 | NA | -30.267 | 139.283 |
| R51794 | Y | N | N | N | NA | NFR | 1998 | NA | -30.198 | 139.233 |
| R52948 | Y | N | N | N | M | NFR | 1999 | NA | -30.121 | 139.399 |

**Table S2.** Sampling sites along a north-south linear transect through the *C. decresii* contact zone. Distance is kilometres (km) from southernmost site 1. Variation (mean ± SD) and samples numbers (*N*) for each trait: hybrid index ($H_{Index}$; Bayesian *q*-value), mtDNA haplotype, quantum catch of the ultraviolet wavelength sensitive receptor ($QC_{UVS}$), PC1 scores for throat coloration ($T_{PC1}$) and dorsolateral phenotype ($D_{PC1}$).

| Ancestry | Site | Dist. | $H_{Index}$ | *N* | mtDNA | *N* | $QC_{UVS}$ | $T_{PC1}$ | *N* | $D_{PC1}$ | *N* |
|---|---|---|---|---|---|---|---|---|---|---|---|
| South | 1 | 0 | 0.98 ± 0.06 | 10 | 1.0 | 10 | 0.65 ± 0.21 | 0.86 ± 0.08 | 22 | 1.65 ± 0.83 | 32 |
| South | 2 | 14 | 0.95 ± 0.13 | 11 | 1.0 | 1 | 0.53 ± 0.14 | 0.76 ± 0.11 | 19 | 1.89 ± 0.90 | 21 |
| Admixed | 3 | 28 | 0.44 ± 0.04 | 11 | 0.18 | 6 | 0.14 ± 0.11 | 0.30 ± 0.22 | 15 | -0.12 ± 0.51 | 18 |
| Admixed | 4 | 31 | 0.40 ± 0.05 | 31 | 0.0 | 10 | 0.22 ± 0.16 | 0.48 ± 0.19 | 37 | -0.19 ± 0.80 | 51 |
| Admixed | 5 | 35.5 | 0.33 ± 0.06 | 10 | 0.0 | 6 | 0.08 ± 0.06 | 0.24 ± 0.21 | 11 | -0.79 ± 0.62 | 11 |
| Admixed | 6 | 39.5 | 0.24 ± 0.04 | 12 | 0.0 | 3 | 0.19 ± 0.12 | 0.43 ± 0.23 | 19 | -0.22 ± 0.60 | 19 |
| Admixed | 7 | 45.5 | 0.16 ± 0.03 | 9 | 0.0 | 1 | 0.09 ± 0.08 | 0.23 ± 0.27 | 6 | -0.39 ± 0.54 | 6 |
| Admixed | 8 | 50.5 | 0.16 ± 0.05 | 7 | 0.0 | 4 | 0.09 ± 0.03 | 0.14 ± 0.01 | 2 | -0.92 ± 0.56 | 2 |
| North | 9 | 58 | 0.09 ± 0.01 | 16 | 0.0 | 5 | 0.18 ± 0.14 | 0.35 ± 0.21 | 24 | -0.59 ± 0.55 | 24 |
| North | 10 | 71 | 0.05 ± 0.01 | 2 | 0.0 | 1 | 0.10 ± 0.04 | 0.46 ± 0.09 | 3 | -0.89 ± 0.58 | 3 |
| North | 11 | 81 | 0.04 ± 0.02 | 9 | 0.0 | 4 | 0.18 ± 0.14 | 0.42 ± 0.12 | 10 | -0.89 ± 0.56 | 10 |
| North | 12 | 94 | 0.02 ± 0.02 | 8 | 0.0 | 9 | 0.28 ± 0.03 | 0.53 ± 0.02 | 4 | -1.18 ± 0.43 | 4 |
| North | 13 | 140 | 0.004 ± 0.02 | 6 | 0.0 | 8 | 0.20 ± 0.12 | 0.35 ± 0.22 | 13 | -0.70 ± 0.57 | 13 |
| North | 14 | 160 | 0.004 ± 0.003 | 7 | 0.0 | 6 | 0.20 ± 0.14 | 0.50 ± 0.18 | 19 | -1.28 ± 0.36 | 19 |
| | | | Total | 149 | - | 74 | - | - | 204 | - | 233 |

**Table S3**. STRUCTURE results summarised in STRUCTUREHARVESTER using the Evanno method for the full dataset of 1333 loci. The selected value of K (number of populations) is denoted with an asterisk.

| K | Reps | Mean LnP(K) | Stdev LnP(K) | Ln'(K) | \|Ln''(K)\| | Delta K |
|---|---|---|---|---|---|---|
| 1 | 10 | -96935.67 | 0.590361 | — | — | — |
| 2* | 10 | -85534.28 | 4.320 | 11401.39 | 8426.593 | 1950.583 |
| 3 | 10 | -82559.483 | 9.957 | 2974.797 | 2381.325 | 164.040 |
| 4 | 10 | -81973.575 | 837.453 | 589.69 | 631.141 | 0.754 |
| 5 | 10 | -80752.744 | 369.785 | 1220.831 | — | — |

**Table S4**. Pairwise $F_{ST}$ values between populations [Northern Flinders Ranges (NFR), Southern Flinders Ranges (SFR), Olary Ranges (OR), Contact Zone (CZ), Mainland South (MS), and Kangaroo Island (KI)]. All values are significant at $P < 0.001$.

| | NFR | SFR | OR | CZ | MS | KI |
|---|---|---|---|---|---|---|
| NFR | - | - | - | - | - | - |
| SFR | 0.173 | - | - | - | - | - |
| OR | 0.211 | 0.027 | - | - | - | - |
| CZ | 0.216 | 0.089 | 0.058 | - | - | - |
| MS | 0.468 | 0.374 | 0.355 | 0.203 | - | - |
| KI | 0.571 | 0.434 | 0.293 | 0.293 | 0.213 | - |

**Table S5.** Likelihoods and parameters from four optimisation rounds for two population demographic models estimated using the program δaδi. Two independent runs were performed for each model. The "sec_contact_asym mig" model, where the species split, followed by a period of isolation before coming into secondary contact with ongoing asymmetrical migration, was the best fit based on log- likelihood (Log-l) and Akaike information criterion (AIC). The parameters were as follows: nu1 = population size of *C. decresii*, nu2 = population size of *C. modestus*, m12 = migration rate from *C. modestus* to *C. decresii*, m21 = migration from *C. decresii* to *C. modestus*, t1 = time between population split and secondary contact, t2 = time between secondary contact and present (sec_contact_asym_mig) or time between secondary contact and isolation (sec_contact_asym_mig_three_epoch), t3 = time between isolation and present.

| **Model** | | **Log-l** | **AIC** | **Chi-squared** | **theta** | **nu1** | **nu2** | **m12** | **m21** | **t1** | **t2** | **t3** |
|---|---|---|---|---|---|---|---|---|---|---|---|---|
| sec_contact_asym_mig | run 1, round 1 | -2970.79 | 5953.58 | 786690.53 | 182.42 | 1.6472 | 4.0666 | 0.68 | 2.01 | 2.1331 | 0.1417 | – |
| | run 1, round 2 | -2882.04 | 5776.08 | 47775.55 | 84.84 | 3.4641 | 8.2668 | 0.2973 | 0.7806 | 6.3964 | 0.3515 | – |
| | run 1, round 3 | -2780.48 | 5572.96 | 1076.29 | 160.81 | 1.6463 | 3.7837 | 0.4455 | 0.6504 | 4.3379 | 0.3209 | – |
| | **run 1, round 4** | **-2666.93** | **5345.86** | **941.79** | **188.33** | **0.9288** | **3.9673** | **0.7834** | **0.7818** | **2.9525** | **0.2833** | – |
| | run 2, round 1 | -2969.25 | 5950.5 | 1792.79 | 360.37 | 0.4698 | 2.0108 | 0.6684 | 0.6881 | 0.8188 | 0.5596 | – |
| | run 2, round 2 | -3026.04 | 6064.08 | 2065.37 | 256.85 | 0.4032 | 3.6384 | 0.8562 | 0.4435 | 1.1506 | 0.3063 | – |
| | run 2, round 3 | -2681.03 | 5374.06 | 1126.49 | 152.28 | 0.8883 | 5.0765 | 0.9495 | 0.8287 | 4.2371 | 0.2464 | – |
| | **run 2, round 4** | **-2672.72** | **5357.44** | **1138.78** | **105.83** | **1.6732** | **7.0967** | **0.4917** | **0.5444** | **6.0399** | **0.3932** | – |
| sec_contact_asym_mig_three_epoch | run 1, round 1 | -3680.41 | 7374.82 | 11090.6 | 96.27 | 3.67 | 6.19 | 0.18 | 0.44 | 4.78 | 2.62 | 0.14 |
| | run 1, round 2 | -3031.38 | 6076.76 | 2265.14 | 84.93 | 2.00 | 10.28 | 0.30 | 0.19 | 4.84 | 1.08 | 0.05 |
| | run 1, round 3 | -3058.94 | 6131.88 | 2398.39 | 99.97 | 0.87 | 9.05 | 0.63 | 0.13 | 4.61 | 1.25 | 0.01 |
| | **run 1, round 4** | **-2869.77** | **5753.54** | **1559.43** | **106.57** | **1.49** | **7.88** | **0.47** | **0.26** | **4.52** | **0.61** | **0.02** |
| | run 2, round 1 | -3936.08 | 7886.16 | 19073.17 | 138.63 | 1.74 | 6.03 | 0.76 | 0.83 | 1.77 | 1.32 | 0.20 |
| | run 2, round 2 | -3530.89 | 7075.78 | 7616.47 | 95.28 | 1.96 | 9.34 | 0.56 | 1.27 | 4.59 | 0.35 | 0.25 |
| | run 2, round 3 | -3054.58 | 6123.16 | 2465.87 | 127.32 | 1.66 | 6.02 | 1.08 | 0.56 | 3.93 | 0.32 | 0.09 |
| | **run 2, round 4** | **-2780.32** | **5574.64** | **1170.49** | **165.73** | **1.24** | **4.19** | **1.05** | **0.90** | **3.83** | **0.21** | **0.02** |

**Table S6**. Results of throat and dorsolateral phenotype analyses on wild-caught males and captive-bred offspring (both sexes for throat and only males for dorsolateral). Contributions of variables to components in principal component analyses (PC1, PC2), linear discriminant analyses (LD1, LD2) analyses, and results of a *t*-test *(T, P*) between northern and southern lineages. Throat variables are the proportions of orange, yellow, and blue colours on the throat, and the quantum catch of the UV receptor ($QC_{UVS}$). Dorsolateral variables are the proportions of orange and yellow colours on dorsal surface, and the number of distinct breaks in the lateral stripe at the neck.

| | Population | Variable | PC1 | PC2 | LD1 | LD2 | *T* | *P* |
|---|---|---|---|---|---|---|---|---|
| Throat | Wild-Caught | Orange | 54.56 | 0.82 | 0.32 | -0.82 | -5.77 | *<0.001* |
| | | Yellow | 9.54 | 69.77 | 0.09 | -0.06 | -1.49 | 0.137 |
| | | Blue | 35.9 | 29.41 | 2.36 | -0.78 | 23.56 | *<0.001* |
| | | $QC_{UVS}$ | - | - | 0.62 | 0.6 | 9.34 | *<0.001* |
| | Captive-Bred | Orange | - | - | -0.09 | -0.5 | -4.26 | *<0.001* |
| | | Yellow | - | - | -0.54 | -0.82 | -5.69 | *<0.001* |
| | | Blue | - | - | 0.59 | -0.82 | 2.27 | *0.04* |
| | | $QC_{UVS}$ | - | - | 1.15 | -0.39 | 8.82 | *<0.001* |
| Dorsolateral | Wild-Caught | Orange | 50.06 | 0.78 | 0.65 | -0.12 | -8.75 | *<0.001* |
| | | Yellow | 0.01 | 98.21 | 0.11 | 1.13 | -3.87 | *<0.001* |
| | | LS Breaks | 49.94 | 1.01 | 1.38 | 0.04 | -30.19 | *<0.001* |
| | Captive-Bred | Orange | - | - | -0.29 | -1.1 | 0.81 | 0.44 |
| | | Yellow | - | - | -0.34 | 0.57 | 2.53 | 0.04 |
| | | LS Breaks | - | - | -1.3 | 0.1 | 19.45 | *<0.001* |

**Table S7.** Parameter estimates for 95% confidence sets (cumulative Akaike weight $w_i \leq 0.95$) of best-ranked cline models fitted using *HZAR* for the hybrid index based on 1333 loci ($H_{index}$), mtDNA haplotype frequencies (mtDNA), PC1 for dorsolateral phenotype ($D_{PC1}$), quantum catch of the UV wavelength sensitive receptor ($QC_{UVS}$), and PC1 for throat coloration ($T_{PC1}$). Cline centre ($c$) is the distance (km) from the southern-most sampling site, width ($w$) is 1/slope, *pmin* and *pmax* are frequencies at cline ends (for genetic loci), $\mu R$ and $\mu L$ are mean trait values at cline ends (for quantitative traits), δ and τ are tail parameters (right, left, or mirror; distance from cline centre to tail and tail slope, respectively). Two log-likelihood unit support limits for centre and width are shown as 'low' and 'high'. Akaike weight ($w_i$) shown has been recalculated based on only the 95% confidence set to sum to 1.

| Trait | Model | $c$ | $c_{low}$ | $c_{high}$ | $w$ | $w_{low}$ | $w_{high}$ | $p_{min}$ / $\mu_R$ | $p_{max}$ / $\mu_L$ | $\delta_L$ | $\tau_L$ | $\delta_R$ | $\tau_R$ | $\delta_M$ | $\tau_M$ | $AIC_c$ | $w_i$ |
|---|---|---|---|---|---|---|---|---|---|---|---|---|---|---|---|---|---|
| $H_{index}$ | *pmin*/*pmax* observed, no tails | 29.47 | 23.96 | 33.92 | 38.55 | 24.60 | 62.00 | 0.00 | 0.98 | - | - | - | - | - | - | 8.18 | 0.23 |
| | *pmin*/*pmax* observed, right tail | 27.74 | 20.09 | 32.12 | 15.65 | 3.45 | 45.15 | 0.00 | 0.98 | - | - | 0.31 | 0.40 | - | - | 8.47 | 0.20 |
| | *pmin*/*pmax* fixed, no tails | 29.00 | 23.16 | 33.66 | 42.59 | 29.15 | 66.34 | 0.00 | 1.00 | - | - | - | - | - | - | 8.82 | 0.17 |
| | *pmin*/*pmax* fixed, right tail | 27.04 | 20.25 | 32.17 | 20.51 | 10.02 | 49.76 | 0.00 | 1.00 | - | - | 0.97 | 0.48 | - | - | 8.93 | 0.16 |
| | *pmin*/*pmax* fixed, mirror tails | 28.78 | 24.15 | 32.93 | 32.07 | 4.90 | 57.43 | 0.00 | 1.00 | - | - | - | - | 14.43 | 0.40 | 10.73 | 0.07 |
| | *pmin*/*pmax* estimated, no tails | 28.32 | 23.60 | 33.32 | 32.44 | 14.77 | 56.92 | 0.03 | 1.00 | - | - | - | - | - | - | 11.07 | 0.05 |
| | *pmin*/*pmax* observed, mirror tails | 29.45 | 24.49 | 33.40 | 31.63 | 2.13 | 57.06 | 0.00 | 0.98 | - | - | - | - | 15.68 | 0.42 | 11.16 | 0.05 |
| | *pmin*/*pmax* observed, left tail | 29.49 | 23.95 | 33.92 | 38.46 | 24.61 | 61.93 | 0.00 | 0.98 | 133.57 | 0.94 | - | - | - | - | 12.38 | 0.03 |
| | *pmin*/*pmax* observed, both tails | 26.46 | 21.51 | 32.17 | 16.58 | 5.11 | 45.45 | 0.00 | 0.98 | 35.23 | 0.44 | 0.31 | 0.42 | - | - | 12.87 | 0.02 |
| | *pmin*/*pmax* estimated, right tail | 27.16 | 22.46 | 31.99 | 20.02 | 7.81 | 46.24 | 0.01 | 0.98 | - | - | 0.22 | 0.51 | - | - | 13.07 | 0.02 |

| | | | | | | | | | | | | | | | | | |
|---|---|---|---|---|---|---|---|---|---|---|---|---|---|---|---|---|---|
| | **Weighted Average** | **28.41** | **22.37** | **33.00** | **29.77** | **15.05** | **55.84** | **0.00** | **0.99** | **4.52** | **0.036** | **0.23** | **0.10** | **1.75** | **0.05** | **-** | **-** |
| mtDNA | *pmin*/*pmax* observed, no tails | 27.70 | 14.75 | 28.17 | 0.80 | 0.02 | 21.88 | 0.00 | 1.00 | - | - | - | - | - | - | 4.17 | 0.29 |
| | *pmin*/*pmax* fixed, no tails | 27.96 | 14.76 | 28.12 | 0.11 | 0.07 | 21.80 | 0.00 | 1.00 | - | - | - | - | - | - | 4.17 | 0.29 |
| | *pmin*/*pmax* fixed, right tail | 27.60 | 14.29 | 28.15 | 1.03 | 0.17 | 21.79 | 0.00 | 1.00 | - | - | 47.72 | 0.81 | - | - | 4.17 | 0.29 |
| | *pmin*/*pmax* observed, right tail | 27.68 | 13.41 | 28.17 | 0.86 | 0.02 | 21.89 | 0.00 | 1.00 | - | - | 103.41 | 0.67 | - | - | 8.58 | 0.03 |
| | *pmin*/*pmax* observed, left tail | 27.94 | 14.74 | 28.17 | 0.16 | 0.01 | 21.88 | 0.00 | 1.00 | 77.57 | 0.71 | - | - | - | - | 8.58 | 0.03 |
| | *pmin*/*pmax* fixed, left tail | 27.76 | 14.81 | 28.14 | 0.62 | 0.05 | 21.86 | 0.00 | 1.00 | 27.18 | 0.59 | - | - | - | - | 8.58 | 0.03 |
| | *pmin*/*pmax* observed, mirror tails | 27.75 | 14.74 | 28.17 | 0.67 | 0.03 | 21.87 | 0.00 | 1.00 | - | - | - | - | 47.82 | 0.28 | 8.58 | 0.03 |
| | **Weighted Average** | **27.78** | **14.59** | **28.17** | **0.64** | **0.08** | **21.85** | **0.00** | **1.0** | **3.35** | **0.04** | **17.20** | **0.26** | **1.53** | **0.01** | **-** | **-** |
| $D_{PC1}$ | mean/variance estimated, right tail | 22.90 | 16.44 | 27.89 | 6.26 | 0.79 | 19.52 | -1.11 | 1.77 | - | - | 0.76 | 0.00 | - | - | 502.71 | 0.77 |
| | mean/variance estimated, mirror tails | 23.43 | 15.96 | 26.70 | 39.49 | 0.92 | 60.43 | -1.66 | 2.89 | - | - | - | - | 9.56 | 0.07 | 505.10 | 0.23 |
| | **Weighted Average** | **23.02** | **16.33** | **27.61** | **13.97** | **0.82** | **29.02** | **-1.24** | **2.03** | **-** | **-** | **0.59** | **0.00** | **2.22** | **0.02** | **-** | **-** |
| $QC_{UVS}$ | mean/variance observed, no tails | 15.85 | 14.3 | 18.93 | 9.04 | 0.88 | 16.49 | 0.20 | 0.65 | - | - | - | - | - | - | -199.63 | 0.71 |
| | mean/variance estimated, no tail | 16.98 | 14.2 | 25.99 | 10.44 | 0.00 | 21.58 | 0.18 | 0.64 | - | - | - | - | - | - | -195.84 | 0.11 |
| | mean/variance observed, left tail | 15.94 | 14.2 | 27.16 | 5.69 | 0.75 | 16.96 | 0.20 | 0.65 | 0.47 | 0.36 | - | - | - | - | -195.63 | 0.10 |
| | mean/variance observed, mirror tails | 14.99 | 14.0 | 19.17 | 4.89 | 0.55 | 18.67 | 0.20 | 0.65 | - | - | - | - | 4.16 | 0.84 | -195.59 | 0.09 |

| | | | | | | | | | | | | | | | | | |
|---|---|---|---|---|---|---|---|---|---|---|---|---|---|---|---|---|---|
| | **Weighted Average** | **15.90** | **14.25** | **20.48** | **8.48** | **0.74** | **17.27** | **0.20** | **0.65** | **0.04** | **0.03** | **-** | **-** | **0.39** | **0.08** | **-** | **-** |
| $T_{PC1}$ | mean/variance estimated, no tails | 15.21 | 15.1 | 16.11 | 2.75 | 2.51 | 6.58 | 0.40 | 0.85 | - | - | - | - | - | - | -96.94 | 0.90 |
| | mean/variance estimated, left tail | 16.94 | 14.7 | 21.00 | 7.40 | 1.78 | 19.96 | 0.39 | 0.86 | 73.72 | 0.95 | - | - | - | - | -92.58 | 0.10 |
| | **Weighted Average** | **15.38** | **15.06** | **16.61** | **3.22** | **2.43** | **7.94** | **0.40** | **0.85** | **7.52** | **0.10** | **-** | **-** | **-** | **-** | **-** | **-** |

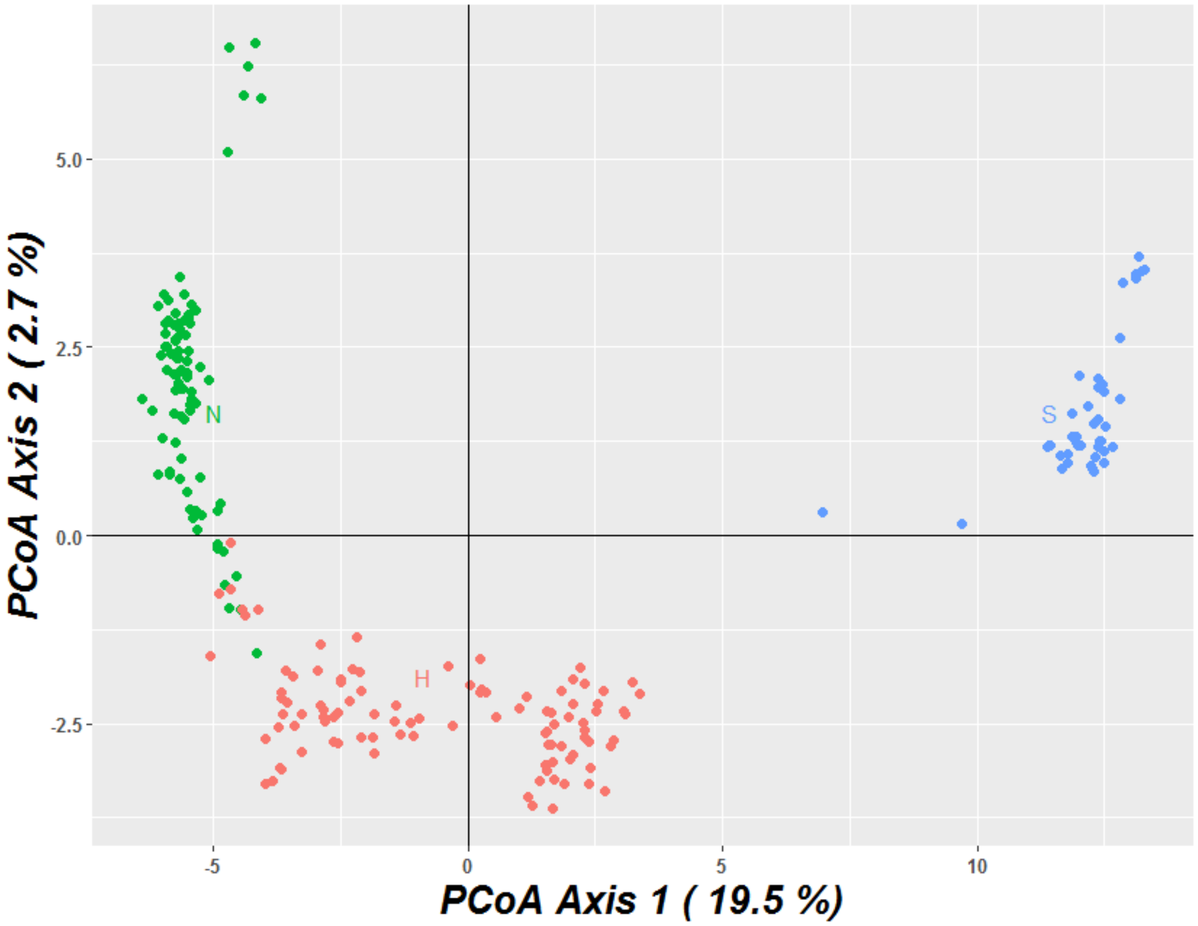


**Figure S1.** Two-dimensional principal coordinate plot (PCoA) constructed from the SNP data set of 6,889 loci showing pairwise genetic distances between individuals: *C. modestus* ("N", green), admixed individuals ("H", red), *C. decresii* ("S", blue).

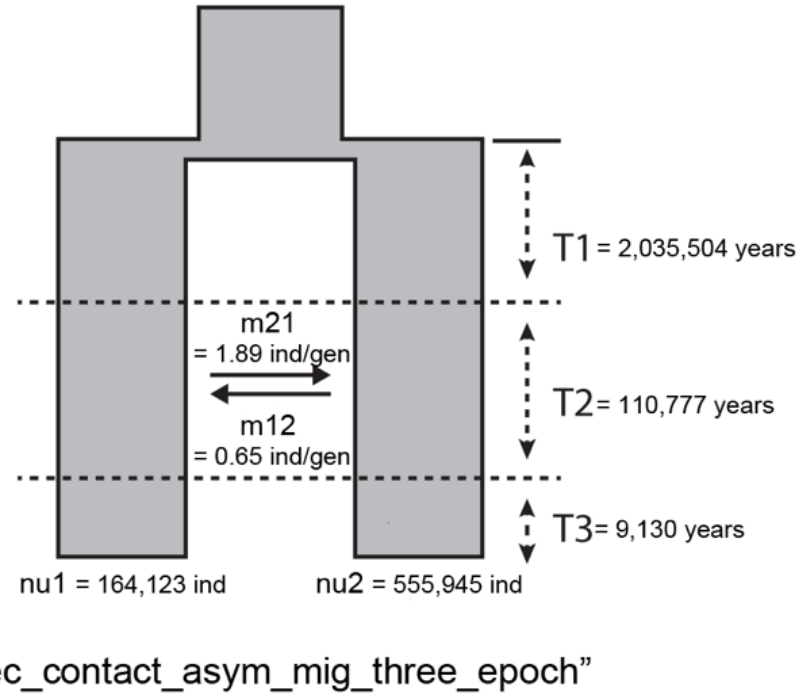


**Figure S2.** Schematic representation of the "sec_contact_asym_mig_three_epoch" demographic model used in δaδi analysis. Populations 1 and 2 are *C. decresii* and *C. modestus* (pooled with admixed individuals) respectively. Converted values are provided for times (T1, T2, T3; in years), migration rates (m12, m21; in individuals per generation) and population sizes (nu1 and nu2; in number of individuals).

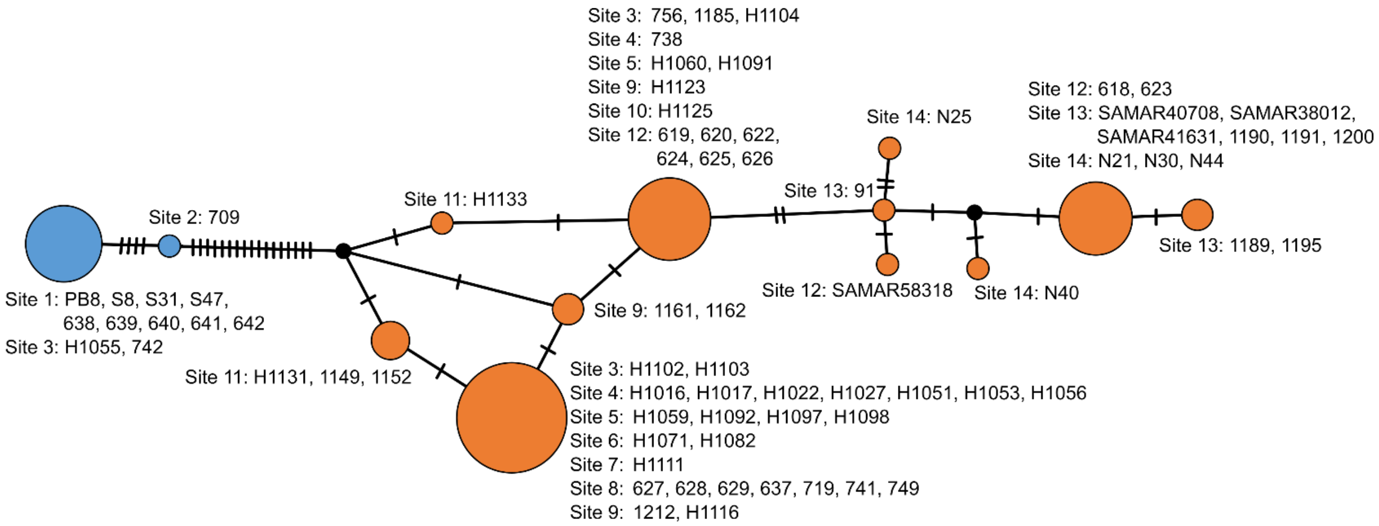


**Figure S3.** TCS parsimony network of *C. decresii* mtDNA haplotypes constructed from the ND4 region. Circle size corresponds to the frequency of haplotypes and each hash indicates one mutational step. Southern lineage haplotypes are coloured blue, northern lineage haplotypes are orange, and hypothetical missing haplotypes are black. Individuals comprising each circle and their geographic origin (cline site) are indicated by adjacent text.

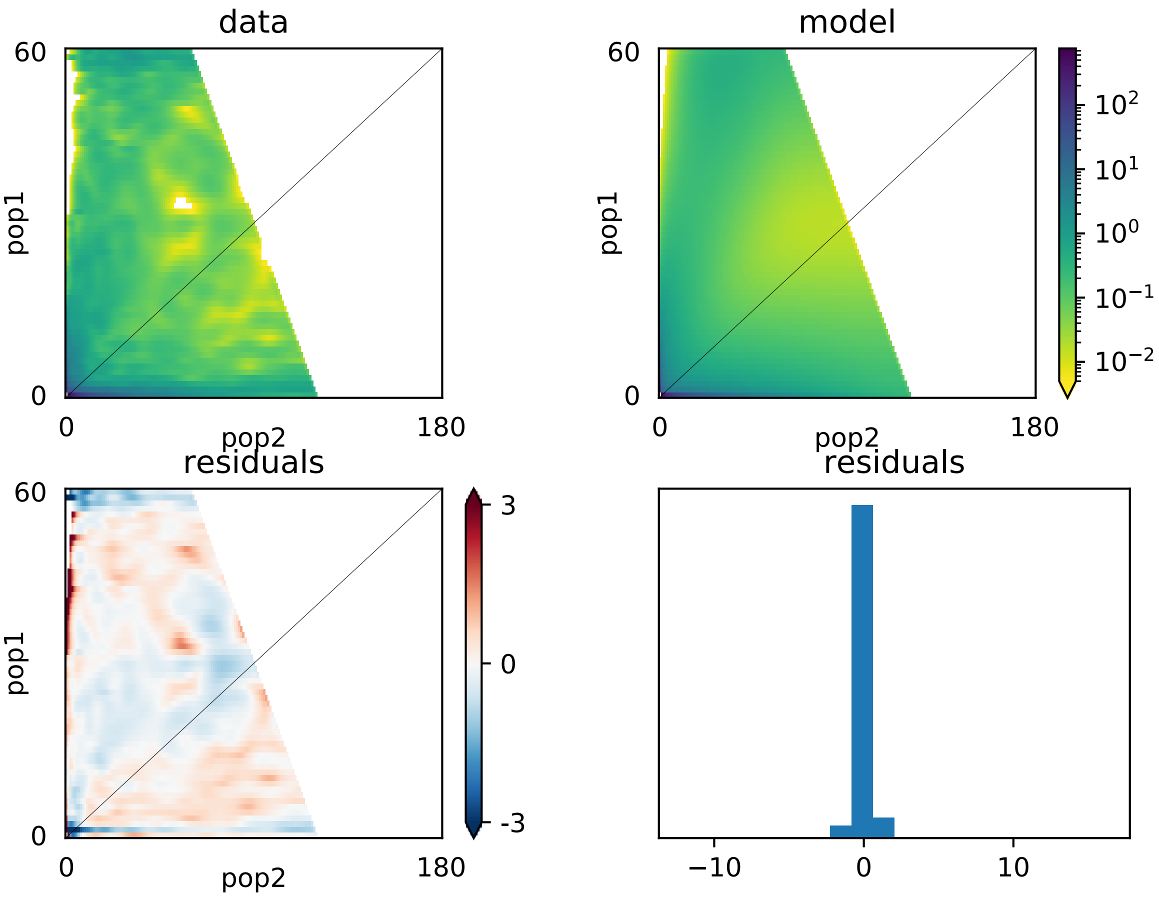


**Figure S4.** The site frequency spectra for the data (top left), and "sec_cont_asym_mig" model (top right), with the location (bottom left), and distribution of model residuals (bottom right). Pop1 is *C. decresii* and pop2 is *C. modestus*.

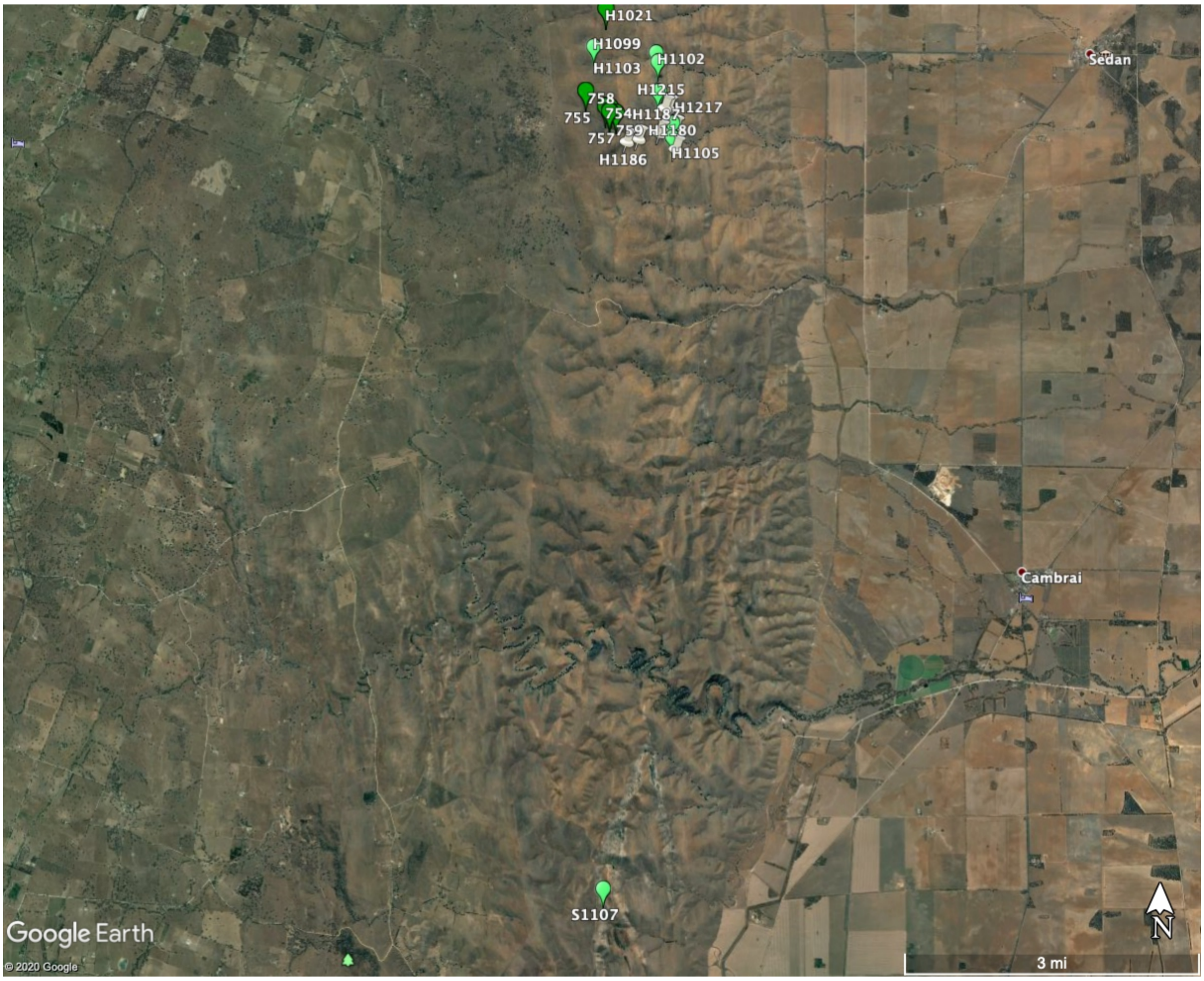


**Figure S5.** Google Earth map showing terrain between site 2 (pure *C. decresii*) and site 3 (admixed).

### Supplemental Materials and Methods

*Captive breeding and offspring rearing*

Adult lizards were housed individually in 55 L x 34 W x 38 D cm opaque plastic enclosures containing a layer of sand and two stacked ceramic tiles for shelter and basking. The room was maintained at temperatures and lighting regimes that mimicked natural seasonal variation. During the breeding season, the ambient temperature was 26°C with a 12:12 light:dark light cycle. A heat lamp was provided in each enclosure to generate a thermal gradient and allow animals to attain their preferred body temperature (36°C; Gibbons, 1977; Walker, unpublished data). Lizards were fed live crickets and mealworms *ad libitum* three times per week and misted with water for hydration.

Our breeding design included four possible combinations (northern female x northern male, northern female x southern male, southern female x southern male, southern female x northern male) with up to three 'rounds' of mating to produce multiple clutches in one breeding season. Female enclosures contained a nest box in the form of a clear plastic container 17 x 13 x 15 cm LxWxD with in a 3.5 cm circular entrance in the lower corner, filled with a moist 50:50 sand:peat moss mixture sloped upwards to the top of the box. Eggs were collected following oviposition and incubated in sealed plastic containers filled two-thirds with moist vermiculite (volume ratio 5:1 vermiculite:water) and incubated at 28°C (± 0.12°C) until hatching for a 50:50 sex ratio (Harlow, 2000). Hatchling lizards were housed individually in 30 x 20 x 10 cm plastic enclosures with fly screen lids. Each enclosure contained two stacked ceramic tiles to provide shelter, paper substrate, water dish, heat lamp and UV lighting. Lizards were fed crickets *ad libitum* and misted with water daily.

*Confirmation of paternity*

Maternity was known for all offspring, however paternity was not known due to multiple mates, and the potential for sperm storage and multiple paternity within clutches in the genus *Ctenophorus* (Hacking, Stuart-Fox, & Gardner, 2017; Lebas, 2001; Olsson, Schwartz, Uller, & Healey, 2007, 2009; Uller, Schwartz, Koglin, & Olsson, 2013). We collected blood from all by venipuncture from the sinus angularis in the corner of the mouth. Genomic DNA was extracted from blood using an E.Z.N.A. Tissue DNA Kit (Omega Bio-tek, Norcross, GA, USA) and sent to the Australian Genome Research Facility (Melbourne, Victoria, AUS) for PCR amplification, fragment visualization and size calling. Paternity was assigned with a 95% confidence level using a maximum likelihood approach in CERVUS v 3.0.7 (Kalinowski, Taper, & Marshall, 2007; Marshall, Slate, Kruuk, & Pemberton, 1998). First, we conducted a simulation of parentage analysis with the parameters of 50,000 offspring, 1% error rate, 5 candidate fathers, 95% of loci typed, and a minimum of 4 loci typed. Following this, we conducted a paternity analysis based on the trio (mother, father, and offspring) LOD score and a strict 95% confidence level (following Rankin, McLean, Kemp, & Stuart-Fox, 2016). These were manually verified across known breeding pairings.

*Testosterone-induced throat colour expression*

Crystalline testosterone powder (no. T1500, Sigma) was mixed with sesame oil at a concentration of 0.025 g/mL oil and 4.5 uL of this mixture was pipetted onto the dorsal surface daily for six weeks until throat coloration was apparent (Rankin et al., 2016). Ventral and dorsal photos and spectral measurements of throat coloration were taken weekly.

*Analysis of phenotypic data*

Photographs were taken using a Canon EOS 600D digital camera, saved in raw format, and included an Xrite colour standard and ruler for scale (Stevens, Párraga, Cuthill, Partridge, & Troscianko, 2007). Images were converted to 8-bit tiff files using Digital Photo Professional (v4.5.0.0; Canon Inc.), linearised with respect to radiance, and equalised against a grey standard (Stevens et al. 2007; as per Smith et al. 2016). We then performed a segmentation analysis (detailed in Teasdale et al., 2013) to extract the proportion of each colour present on the throat (i.e. orange, yellow, blue), classify throats into existing morphs (orange, yellow, orange-yellow, grey, or blue), and to quantify the proportion of orange and yellow colouration present on the head and neck from dorsal photographs. Image analysis was conducted using custom code written in MATLAB (The MathWorks, Inc., Natick, MA, USA; see Teasdale et al., 2013).

We took spectral reflectance measurements (300 – 700 nm) to quantify variation in UV reflectance, using an Ocean Optics (Dunedin, FL, USA) Jaz portable spectrometer. The probe was held at a constant angle (45°) and distance from the surface of the skin. Measurements were expressed relative to a Spectralon 99% diffuse white reflectance standard (Labsphere, Inc., North Sutton, NH, USA). At least two measurements were taken of each colour present on the throat, then averaged. For each throat colour patch, we estimated the receptor quantum catch (QC), the stimulation of retinal photoreceptors, for the violet to UV wavelength sensitive (UVS) photoreceptor because we were specifically interested in the diagnostic UV-blue component of throat coloration. The R package pavo (v1.3.1; Maia, Gruson, Endler, & White, 2018) was used to process reflectance measurements.
